# Characterization of the NiRAN domain from RNA-dependent RNA polymerase provides insights into a potential therapeutic target against SARS-CoV-2

**DOI:** 10.1101/2021.02.03.429510

**Authors:** Abhisek Dwivedy, Richard Mariadasse, Mohammed Ahmad, Sayan Chakraborty, Deepsikha Kar, Satish Tiwari, Tanmay Majumdar, Jeyaraman Jeyakanthan, Bichitra K. Biswal

## Abstract

Apart from the canonical fingers, palm and thumb domains, the RNA dependent RNA polymerases (RdRp) from the viral order *Nidovirales* possess two additional domains. Of these, the function of the Nidovirus RdRp associated nucleotidyl transferase domain (NiRAN) remains unanswered. The elucidation of the 3D structure of RdRp from the severe acute respiratory syndrome coronavirus-2 (SARS-CoV-2), provided the first ever insights into the domain organisation and possible functional characteristics of the NiRAN domain. Using *in silico* tools, we predict that the NiRAN domain assumes a kinase or phosphotransferase like fold and binds nucleoside triphosphates at its proposed active site. Additionally, using molecular docking we have predicted the binding of three widely used kinase inhibitors and five well characterized anti-microbial compounds at the NiRAN domain active site along with their drug-likeliness as well as DFT properties. For the first time ever, using basic biochemical tools, this study shows the presence of a kinase like activity exhibited by the SARS-CoV-2 RdRp. Interestingly, the proposed kinase inhibitors and a few of the predicted nucleotidyl transferase inhibitors significantly inhibited the aforementioned enzymatic activity. In line with the current global COVID-19 pandemic urgency and the emergence of newer strains with significantly higher infectivity, this study provides a new anti-SARS-CoV-2 drug target and potential lead compounds for drug repurposing against SARS-CoV-2.

## Introduction

RNA dependent RNA polymerases (RdRp) are conserved across all RNA virus superfamilies harbouring a positive or negative sense RNA genome with the exception of retroviruses. RdRp catalyses the replication of RNA genomes from an RNA template, thus plays a key role in the viral life cycle and pathogenesis [1]. The structure of RdRp is broadly divided into three domains-fingers, palm and thumb, the shape resembling that of a right hand. This structural organization is conserved not only in RNA viruses, but also in the DNA polymerases across all kingdoms of life [1,2]. However, the positive-stranded RNA (+RNA) viruses from the order *Nidovirales* present a conspicuous difference in the structural organization of their RdRp molecules [2–4]. The viruses in the order *Nidovirales* are known to infect a broad spectrum of hosts. While members of the families *Arteriviridae* and *Coronaviridae* infect vertebrates, members of *Mesoniviridae* and *Roniviridae* primarily infect invertebrates [2–4]. Despite a significant difference in the sizes of their genomes which ranges from 13 kb to 35 kb, the RdRp molecules encoded by each genome show a common structural organization [2].

Apart from the basic RdRp structural organisation comprised of the canonical palm, thumb and finger domains, the Nidovirus RdRp possesses two additional domains-the Nidovirus RdRp associated nucleotidyl transferase domain (NiRAN) and the interface domain [2–6]. The interface domain primarily serves as a connector between the NiRAN and canonical RdRp regions. However information on the functional aspects of the NiRAN domain remains limited. The first ever study that defines the NiRAN domain from the RdRp/Nsp9 of equine arteritis virus (EAV), demonstrates a manganese-dependent covalent binding of guanosine phosphate and uridine phosphate to an invariant lysine residue from the NiRAN domain [3]. The study proposes three possible molecular functions of the NiRAN domain: as a ligase, as a GTP dependent 5’ nucleotidyl transferase and as a UTP dependent protein priming function facilitating the initiation of RNA replication [2,3]. Various studies from other RNA viruses have demonstrated the nucleotidyl transferase activities exhibited by the N-terminal regions of the respective RdRp molecules [7–9]. However these transferase activities have been attributed to both 5’-priming as well as terminal ribonucleotide addition functions. Moreover, RdRp enzymes are known to exhibit either a primer dependent or a primer independent initiation of RNA replication [1,10]. Interestingly, nidovirus RdRp molecules have been experimentally demonstrated to possess both modes of initiation, thus hinting as a probable UTP mediated priming function of the NiRAN domain [11,12].

An earlier cryo-EM structure of replicase polyprotein complex from SARS-CoV had significant regions of the NiRAN domain missing, thus failed to provide any functional association to this domain [5]. However, a recent study reporting the cryo-EM structure of the replicase polyprotein complex from SARS-CoV-2 provides the complete structural preview of the NiRAN domain [6]. Together, these studies suggest that the interface domain also serves as the binding partner for Nsp8 protein of the polyprotein complex. COVID-19, the disease caused by the nidovirus SARS-CoV-2 is an on-going global crisis that necessitates the exploration of viral protein functions in addition to the exploration for novel anti-coronavirus inhibitor scaffolds [13–15]. In addition, the recent emergence of newer SARS-CoV-2 strains with significantly higher contagion [16] further necessitates the discovery of novel anti-viral targets and small molecule inhibitors targeting the life cycle and metabolism of the virus. In this study, utilizing a combination of *in silico* and biochemical tools, we propose that the NiRAN domain of SARS-CoV-2 possess a kinase like fold and is possibly involved in the catalysis of a kinase/phosphotransferase reaction. In addition, we also report the structure aided docking of a number of known kinases and nucleotidyl transferase inhibitors and their inhibitory potential on the kinase like enzymatic activity of SARS-CoV-2 RdRp.

## Results

### Analysis of the NiRAN and Interface domains

As described in a recent study [6], the NiRAN and the interface domains span over residues 1-365 of the SARS-CoV-2 RdRp polypeptide sequence. These two domains form an arrow head like structure, which acts as a base for the RdRp region of protein (Fig. 1A). Of these, amino acids 4-28 and 69-249 have been designated as the NiRAN domain that comprises of 8 α-helices (H1-H8) and 5-β strands (S1-S5), respectively. Two independent β-strands (B1, B2) form a hairpin like structure (residues 29-68) in the vicinity of NiRAN domain and are not considered integral to the NiRAN domain. The interface domain stretches from residue 250 to residue 365 and is comprised of a 6 α-helix bundle (h1-h6) followed by 3 antiparallel β strands (s1-s3). The overall topology suggests that the two distinct domains are connected by a small linker of 1 residue (Fig. 1B). As proposed in the EAV-NiRAN study, several key residues such as K94, R124, S129, D132, D165 and F166 [3], were found conserved in the SARS-CoV-2-NiRAN as revealed in a pairwise alignment of the two sequences. The corresponding residues from SARS-CoV-2-NiRAN are K73, R116, T123, D126, D218 and F219. These residues have been predicted to be crucial for the enzymatic function of the NiRAN domain. A multiple sequence alignment of 17 RdRp polypeptide sequences from coronaviruses across multiple host species (human, bat, cow, pigs and rodents) revealed an absolute conservation of these residues (Fig. S1). Apart from the aforementioned residues, 78 (total of 84; 45 in NiRAN and 39 in interface domains) other residues were found strictly conserved in the RdRp molecules across all 17 viruses, hinting at a key role of these residues in the enzymatic and/or functional properties of NiRAN and Interface domains. A visualization of the conserved residues of the NiRAN domain shows that a significant majority of these residues are localized between the 4 stranded β-sheet and the following helix bundle (Fig. 1B). A surface potential representation of the regions reveals the presence of five potential entry pockets suggesting that the active site of the NiRAN domain is possibly flanked by these conserved residues (Fig. S2A). The 39 residues conserved in the interface domain are however spread all over the domain architecture (Fig. S2B). The conservation of the residues across species suggests an important role of the NiRAN domain in RdRp mediated RNA replication in coronavirus, as suggested by the previous study on EAV-RdRp [3].

**Figure 1:**
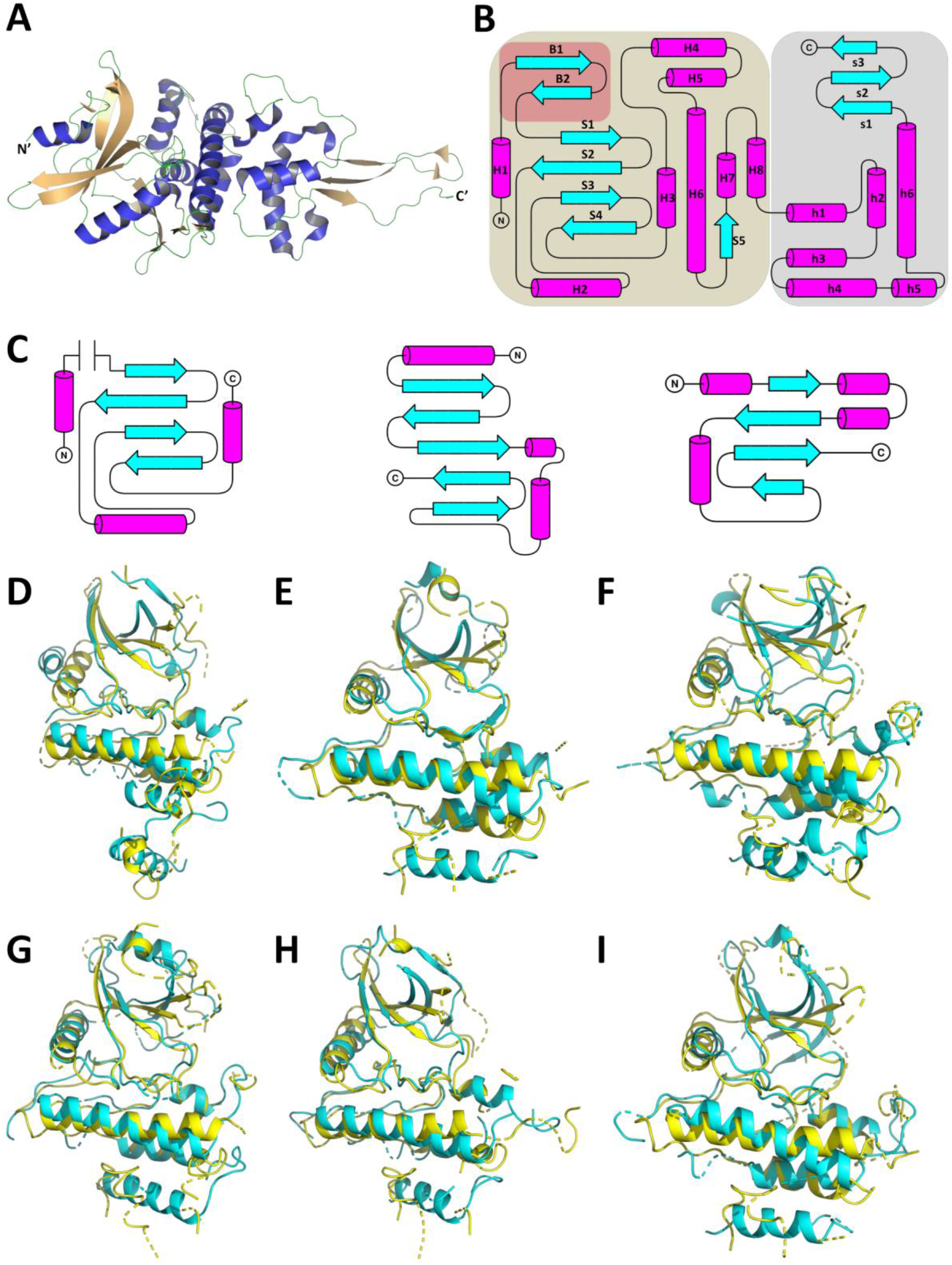
The CoV-2-RdRp NiRAN domain present a kinase/phosphotransferase like structural organisation. **(A)** The NiRAN and the interface domains of the CoV-2-RdRp present an arrow head like structure (helices in royal blue, strands in peach and loops in green). **(B)** The overall topology of the NiRAN (brown background, helices as marked as H and strand are marked as S) and interface domain (grey background, helices as marked as h and strand are marked as s) along with β-hairpin structure (brick red background, strand are marked as B). **(C)** The NiRAN domain possess a topology (left panel) that borrows elements from the canonical kinase fold (centre panel) as well as the non-canonical kinase fold of TgBPK1 (right panel). **(D-I)** The structural superimpositions of the secondary structural elements of CoV-2-RdRp NiRAN with known kinases reveal significant alignment in the antiparallel β-sheet and in the helices that follow. **(D)** Lim 2 kinase domain. **(E)** Syk kinase domain **(F)** O-mannosyl kinase domain. **(G)** IRAK4 kinase domain. **(H)** FGFR2 kinase domain. **(I)** Insulin receptor kinase domain. (The aforementioned kinases’ structural elements are shown in cyan, CoV-2-RdRp NiRAN structural elements shown in yellow.)

### Prediction of the plausible functions of the NiRAN domain

The understanding of the overall topology, localization of the conserved residues prompted an investigation on understanding the enzymatic function of the NiRAN domain. For this, two different approaches were considered with the Protein data bank as the target database. The first approach utilized the prediction of Hidden Markov Models (HHPred) [17] while the second approach predicted the presence of similar folds (ORION) [18]. The list of the top 15 hits retrieved for the first and second approaches are listed in Tables S1 and S2, respectively. Notably, both approaches predicted a variety of kinase and kinase like transferase (phosphotransferase molecules). A previous report had evidenced at the similarity of the NiRAN domain to that of a known pseudokinase molecule SelO [5]; however further characterization was not feasible due to absence of the NiRAN 3D structure. We compared the topologies of the NiRAN domain with that of the canonical kinase fold. Similar to the canonical kinase fold, the NiRAN domain comprises of an antiparallel β-sheet flanked by alpha helices. The canonical kinase domain exhibits a 5 stranded antiparallel β-sheet, which is flanked by two helices running parallel and one helix running perpendicular to the β-sheet (Fig. 1C, middle panel) [19,20]. However, the NiRAN domain shows a 4 stranded antiparallel β-sheet (S1-S4) flanked by one parallel (H2) and two perpendicular helices (H1 and H3) (Fig. 1C, left panel). Further investigation into the available literature confirmed the presence of many non-canonical kinase folds, one of which presents a 4 stranded antiparallel β-sheet flanked by three parallel and one perpendicular helix (Fig. 1C, left panel) [21]. Taken together, these results suggest that the NiRAN domain assumes a kinase like fold, possibly functioning either as a pseudokinase or a phosphotransferase.

To further investigate this proposition, a structural alignment of the NiRAN domain was performed against 8 randomly selected kinase molecules retrieved in the HHPred and ORION searches. The molecules Lim 2 kinase domain (PDB ID-5NXD) (Fig. 1D, Fig. S3A and S4A), Syk kinase domain (PDB ID-4YJR) (Fig. 1E, Fig. S3B and S4B), O-mannosyl kinase domain (PDB ID-5GZ9) (Fig. 1F, Fig. S3C and S4C) and IRAK4 kinase domain (PDB ID-2NRU) (Fig. 1G, Fig. S3D and S4D) aligned with the NiRAN domain with rmsd values ranging from 0.6 - 4 Å and with alignment scores ≥ 0.5, suggesting that the kinase domains in these molecules share nearly similar fold with the NiRAN domain. As a proof of principle, two additional kinase molecules not listed in the aforementioned search results-FGFR2 kinase domain (PDB ID-2PVF) (Fig. 1H, Fig. S3E and S4E); and Insulin receptor kinase domain (PDB ID-1GAG) (Fig. 1I, Fig. S3F and S4F) were aligned with the NiRAN domain. Both these molecules displayed rmsd values ranging from 0.22 - 4 Å and alignment score of ∼0.5, further highlighting the possible presence of a kinase like fold in the NiRAN domain. The results clearly exhibited the alignment of both the Cα backbone of the two molecules and the conservation of secondary structural elements. Of note, the alignments of the helix H2, helix H6, helix H8 and the antiparallel β-sheet (S1-S4) are strikingly evident. Of note, the previously mentioned pseudokinase molecule SelO (PDB ID-6EAC) also aligned with the NiRAN domain. With rmsd values ranging from 2 - 8 Å and an alignment score of 0.42, the alignment suggests that though both the molecules roughly assume similar folds, the amino acid conservation differs significantly. Collectively, results of the alignments further highlight the presence of a non-canonical kinase/phosphotransferase like fold within the NiRAN domain.

### Predicting the NiRAN domain active site

The earlier study with EAV-RdRp experimentally demonstrated the binding of GTP and UTP nucleotides to the NiRAN domain [3]. A docking analysis of these nucleotides with the NiRAN domain was performed to delineate the probable active site. Kinase domains in general possess an active site between the antiparallel β-sheet and the subsequent helix bundle [19,22,23]. Both GTP (Fig. 2 A & A’) and UTP (Fig. 2 B & B’) docked well within the probable active site region with docking scores of -9.84 kcal/mol and -6.59 kcal/mol, respectively. Besides, end-point binding free energy calculation with MM/GBSA suggest a strong interaction between the nucleotides and the residues in the probable active site (-15.77 kcal/mol and -17.67 kcal/mol for GTP and UTP, respectively). Examination of the molecular interactions suggests that the GTP molecule formed possible H-bonding with R116 and salt bridge interaction with K73. The UTP molecule however only displayed a salt bridge interaction with K73. It is worth mentioning that mutation of the corresponding K94 residue in EAV-RdRp resulted in complete loss of nucleotidylation activity for both GTP and UTP [3]. Using the above interactions and the critically conserved residues, we computationally predicted the active site of NiRAN domain. The predicted active site pocket is lined with the following residues: F35, D36, I37, Y38, N39, F48, L49, K50, T51, N52, R55, V71, <u>K73</u>, R74, H75, N79, <u>E83</u>, <u>R116</u>, <u>L119</u>, <u>T120</u>, K121, Y122, <u>T123</u>, V204, <u>T206</u>, <u>N208</u>, <u>N209</u>, <u>Y217</u>, <u>D218</u>, <u>G220</u>, <u>D221</u>, F222 and <u>S236</u>; the underlined residues being strictly conserved across coronavirus RdRp molecules. K73 has been predicted as one of the key residues in the NiRAN active site. The corresponding lysine from the EAV-RdRp (K94) has been deemed critical to the function of NiRAN domain, owing to its interactions with GTP and UTP. The study also demonstrated that mutating lysine to alanine results in complete loss of RdRp function [3]. An overview of the active site pocket depicting the electrostatic surface potential indicates the localization of charged residues at the entry points to the site, while the interior of the pocket remains lined with largely non polar residues (Fig. 2C).

**Figure 2:**
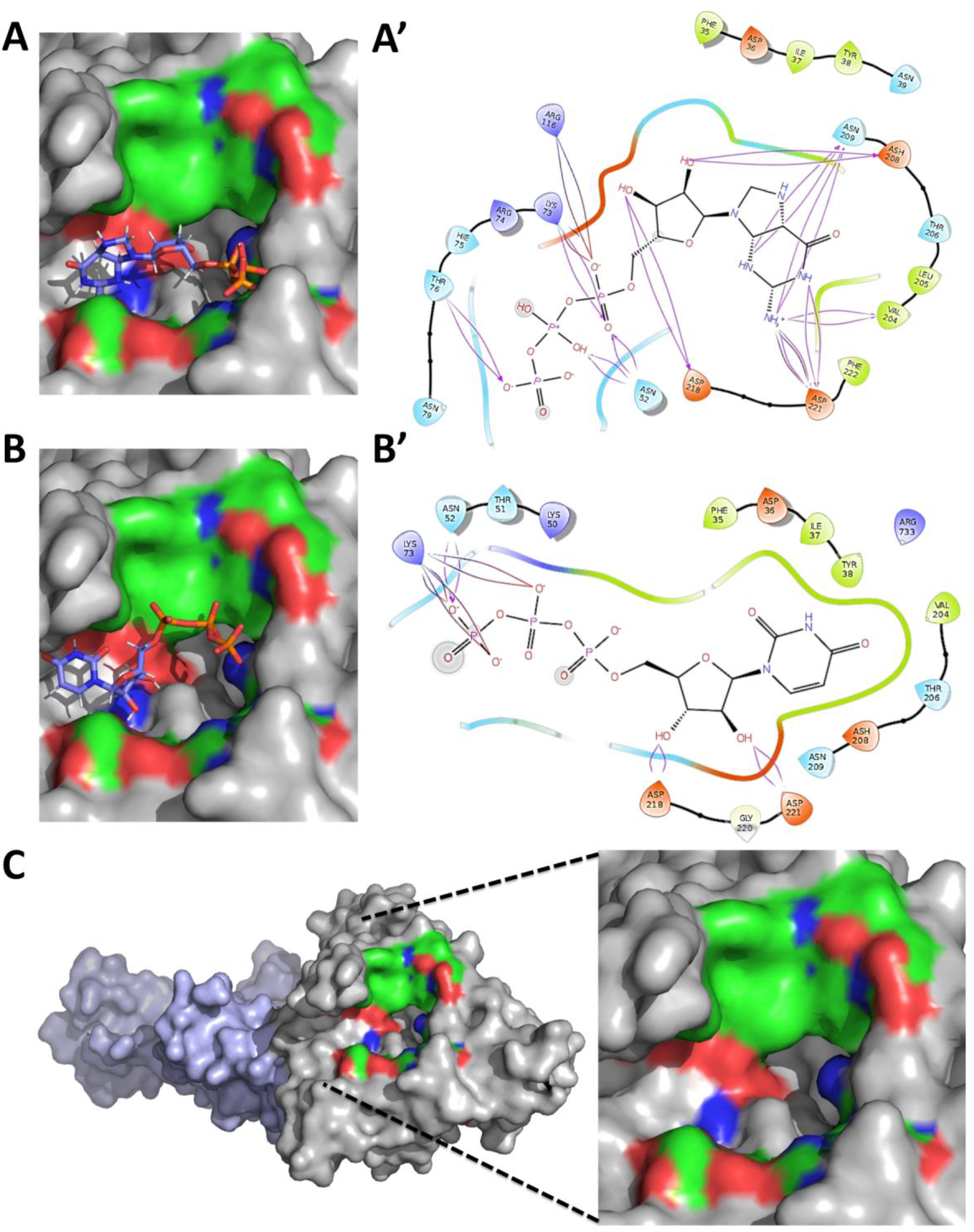
The active site of CoV-2-RdRp NiRAN domain binds GTP and UTP and exhibits kinase like motifs. GTP and UTP bind at the probable active site of the NiRAN domain with notably low free binding energies. **(A)** GTP within the active site pocket. **(B)** UTP within the active site pocket. **(C)** GTP binding reveals salt bridge interaction with K73 and H-bonding with Asp 116. **(D)** UTP binding exhibits salt bridge interaction with K73. **(E)** The predicted active site of the NiRAN domain possess both charged and uncharged residues, where in the charged residues primarily line the entry points of the pocket and the uncharged residues line in the deeper sections. (Red indicates positively charged regions, blue indicates negatively charged regions and green indicates neutral regions, grey indicates regions beyond GTP-binding pocket).

With the above results suggesting a kinase like fold and the active site showing possible GTP and UTP binding function, a motif search was performed using the complete sequence of the NiRAN, β-hairpin and interface domains. The results predicted the presence of kinase like motifs belonging to Protein kinase C family, Protein kinase A family and Src Kinase family [24]. Few sequences motifs belonging to unspecified kinase families were also predicted. Interestingly, no kinase like motif was predicted for the interface domain and the β-hairpin region. Of note, a myristoylation consensus sequence overlapping with an unspecified kinase consensus was also predicted. Host enzymes are known to interact with viral proteins inducing post translational modifications following host or virus mediated signals. In particular, host kinases are known to post translationally modify viral protein often resulting in enhanced virulence or induction of various metabolic pathways [25–27]. A kinase site prediction suggested 6 different kinase phosphorylation sites with specificity for the eIF2 kinase family. A sequence based visualization of these motifs is presented in Fig. S5. These predictions hint at possible post translational modifications of viral proteins, mediated by the interplay of host – SARS-CoV-2, possibly aimed at regulating the host and viral metabolism.

In order to further explore the kinase like catalytic nature of NiRAN domain, three broad specificity kinase inhibitors-Sunitinib, Sorafenib and SU6656 were randomly selected and docked into the predicted active site of the NiRAN domain. All three kinase inhibitors show strong binding at the predicted active site as evident from the low free energy of binding (Fig. 3 A, B and C). Interestingly, these inhibitors also demonstrate potential H-bond with residues lining the active site. While Sunitinib and Sorafenib primarily interact with aspartate residues, SU6656 exhibits H-bond interaction with lysine residue 73 (Fig. 3 A’, B’ and C’), which is the critical lysine residue predicted to be involved in NiRAN domain’s catalytic activity, as mentioned earlier. The results of the docking analysis are presented in Table S3. In a nutshell, the above predictions further suggest that the NiRAN domain of coronavirus RdRp has a kinase/phosphotransferase like catalytic activity.

**Figure 3:**
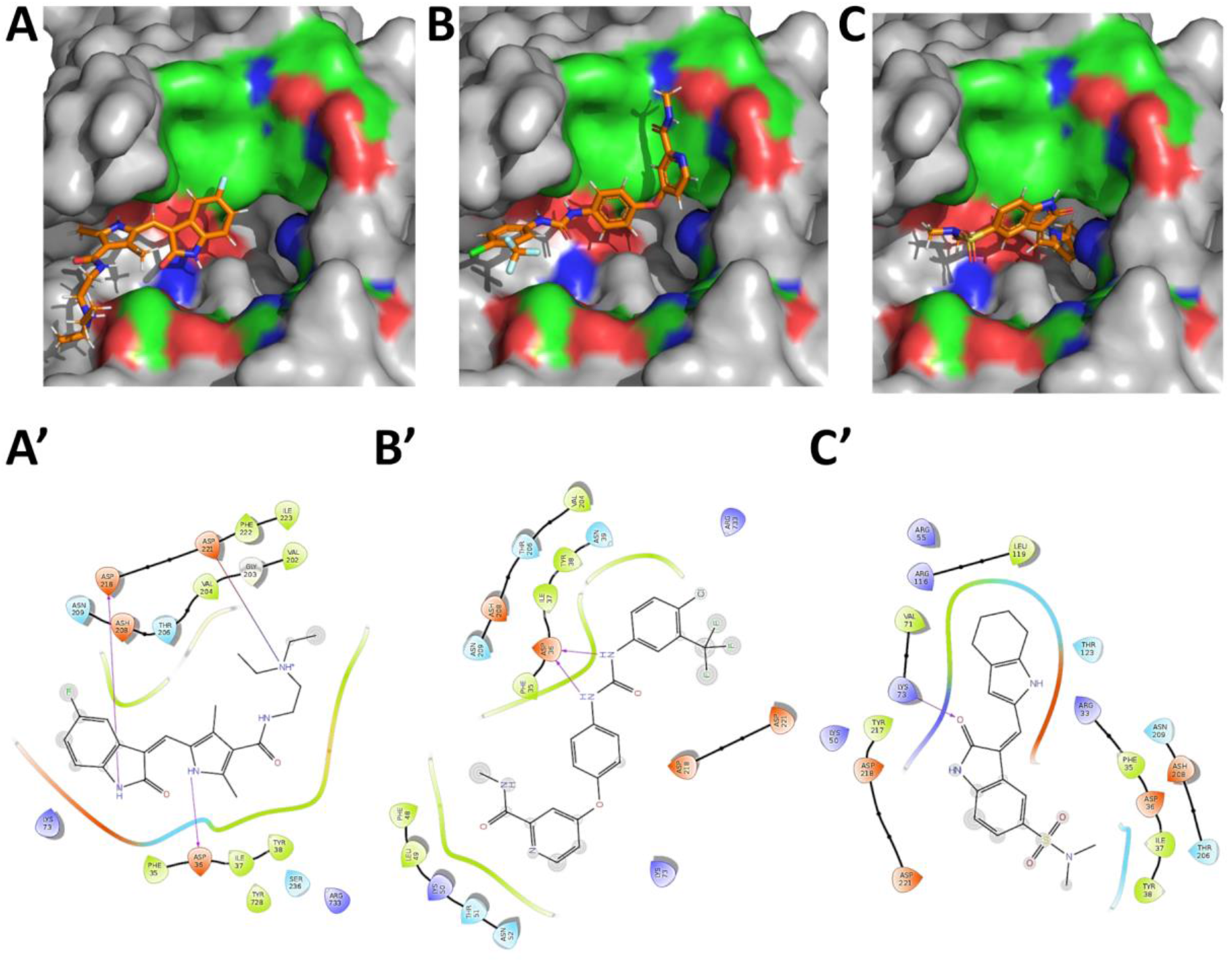
The active site of CoV-2-RdRp NiRAN domain binds kinase inhibitors. **(A, C and E)** Broad specificity kinase inhibitors bind within the predicted NiRAN active site with significantly low free binding energies. (Red indicates positively charged regions, blue indicates negatively charged regions and green indicates neutral regions, grey indicates regions beyond GTP-binding pocket): **(A)** Sunitinib within the active site pocket; **(B**) Sorafenib within the active site pocket; and **(C)** SU6656 within the active site pocket. The kinases inhibitors demonstrate H-bond interactions between with the enzymatically critical aspartate and lysine residues lining the active site-**(A’)** Sunitinib; **(B’)** Sorafenib; and **(C’)** SU6656.

### Prediction of inhibitors targeting the NiRAN domain

As mentioned earlier, this domain might be involved in GTP induced protein phosphorylation, thus enabling a primer independent RNA replication (G-capping) [3]. Also, the domain may be involved in phosphorylation of UTP, thus functioning as a terminal nucleotidyl transferase. A wide range of viruses, both with DNA and RNA genomes are known to possess either multifunctional or dedicated proteins exhibiting the aforementioned activities [7,28]. Interestingly, many pathogenic bacteria also encode for proteins that possess G-capping and/or terminal nucleotidyl transferase activities [29,30]. Years of research have also presented a plethora of inhibitor molecules that target these essential functions in pathogens that cause diseases in human and non-human mammals [28]. After careful examination, a list of 77 compounds with experimentally demonstrated inhibitory potential against members of *Flaviviridae, Togaviridae*; Human cytomegalovirus, Herpes simplex virus, and few gram negative bacterial species were selected for docking against the CoV-2-RdRp NiRAN domain active site [28,31–37]. The list of the selected compounds is presented in Table S4.

Further, this study will discuss the best five inhibitors based on the docking analysis performed on the aforementioned 77 compounds. The five inhibitors with the best docking in scores in a decreasing order are-65482, 122108, 135659024, 4534, and 23673624 (numbers represent the PubChem IDs). The results of the docking analysis are presented in Table S5. All these five inhibitors occupy varying regions within the active site pocket in a manner that their aromatic rings align with the uncharged/non-polar regions, while the charged moieties fit in the extremities of the active site pocket (Fig. 4 A-E). All the five inhibitors exhibit H-bonding interactions with the active site residues. Interestingly all the five inhibitors present potential salt bridges with active site residues and also exhibit interactions with enzymatically critical residue K73, either via H-bonding or salt bridges. In addition to the above interactions, Compound 65482 also exhibits the pi-pi interactions between its own purine ring and residues F35 and F48, as is evident from its best docking score. The molecular interactions of these inhibitors with the CoV-2-RdRp NiRAN domain active site are presented in Fig. 6 A’-E’. In order to estimate the drug-likeliness of the aforementioned inhibitors, an ADME/T analysis was performed. The ADME/T analysis failed to provide any results for the compound 135659024. The ADME/T properties of the other four molecules were within the acceptable range in terms of Lipinski’s rule of five [38] and related pharmacokinetic criteria such as H-bonding atoms probability, human oral absorption and the IC_50_ values for blockage of human Ether-a-go-go-Related Gene (hERG) K^+^. The results of this analysis are listed in Table S6.

**Figure 4:**
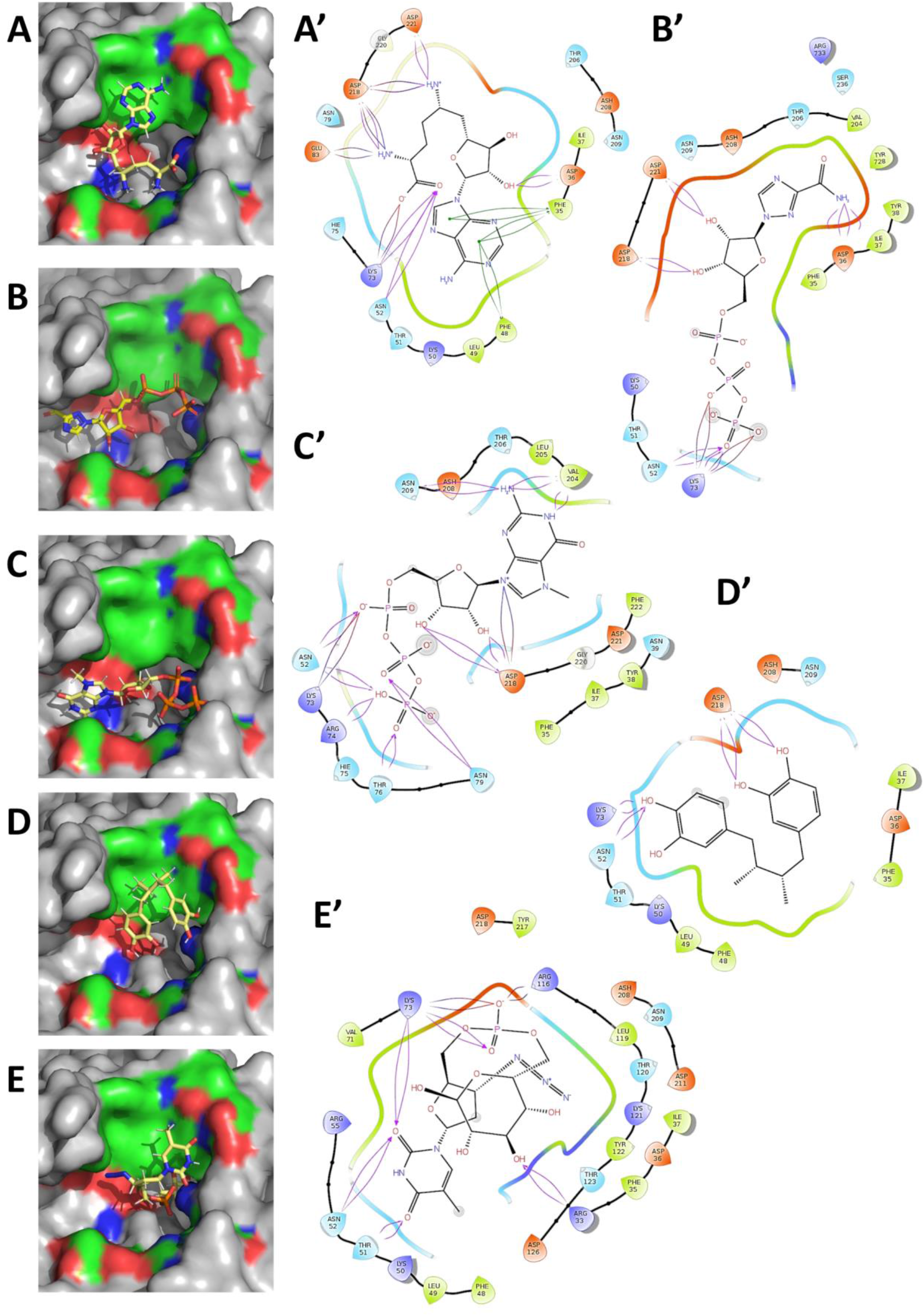
The binding and molecular interaction of the best five predicted nucleotidyl transferase inhibitors at the active site of CoV-2-RdRp NiRAN. The computationally directed binding of-**(A)** 65482; **(B)** 122108; **(C)** 135659024; **(D)** 4534; and **(E)** 23673624; within the active site pocket. (Red indicates positively charged regions, blue indicates negatively charged regions and green indicates neutral regions, grey indicates regions beyond GTP-binding pocket). The molecular interactions between the inhibitors and the active site pocket reveal H-bonds, salt bridges and pi-pi interactions-**(A’)** 65482; **(B’)** 122108; **(C’)** 135659024; **(D’)** 4534; **(E’)** 23673624. Of note, the best predicted inhibitor-65482 presents all the aforementioned molecular interactions.

### Molecular electronic features of predicted inhibitors

The quantum mechanical features of the best five inhibitors were investigated to understand the chemical reactivity of the molecules against the NiRAN active site. The Highest Occupied Molecular Orbital (HOMO) and Lowest Unoccupied Molecular Orbital (LUMO) of five inhibitors describe the electron donate (electron rich) and accepting (low electrons) regions respectively. Molecular electrostatic potential energy defines the positive and negative charge distribution of the molecule for necessary reactivity. In terms of HOMO and LUMO, it is pertinent to understand the molecular properties of the inhibitors; in order to better predict and comprehend the electron transfer reaction (nucleophilic and electrophilic attack). Localization of HOMO, LUMO and electrostatic potential surface density of five molecules are presented in Fig. S6 A-E.

In the inhibitor 65482, both the HOMO and LUMO occupy regions in the adenosine ring. HOMO occupies significantly more regions, thus suggesting a possible nucleophilic reaction by the inhibitor. In term of electrostatic potential surface, the inhibitor 65482 consists of highly negatively charged surface around the molecule, the only positive charged surface being around the hydroxyl group of the side chain. In the inhibitor 122108, the HOMO region localizes in the phosphate group while the LUMO regions encompasses the triazole-3-carboxamide group. Also, a strong positive charged surface is present around the phosphate group. In the inhibitor 135659024, HOMO and LUMO regions cover the adenosine ring and a highly positive surface charge density is located around the phosphate group, thus indicative of a possible nucleophilic reaction. In case of the inhibitor 4534, HOMO regions localize in both the catechol rings, while LUMO encompasses a single catechol ring of the molecule. With a significantly higher proportion of HOMO regions is indicative a nucleophilic reaction. Also, the negative charge surface potential is localized in small region in one of the two catechol rings, with positive charge surface potential encompassing the remain regions of the inhibitor. For the inhibitor 23673624, the HOMO regions occupy the pyrimidine ring and the LUMO regions occupy the azide anion group. In term of electrostatic surface potential, a high negative charge is spread over the molecule while the positive surface charge covers the phosphate group. The energy values and HOMO-LUMO energy gap of all the five inhibitors are presented in Table S7. Lowering the energy gap between HOMO-LUMO couples is known to improve the chemical reactivity of the inhibitors.

### The predicted molecules inhibit a kinase like activity exhibited by the NiRAN domain

In order to determine any putative kinase like activity being harboured by the CoV-2 RdRp, the protein was overexpressed, purified (Figure S7 A & B) and its identity was confirmed by mass spectrometry (Figure S7C & Table S8). The absence of any known kinase/phosphotransferase substrate for CoV2-RdRp posed a different challenge. Most kinases are specific and rarely phosphorylate other targets [39]. In addition, in the absence of any substrate, kinases have been shown to exhibit a residual intrinsic ATPase/GTPase like activity [40–43]. Here, the enzyme enzymatically cleaves the nucleoside triphosphate molecule and transfers the gamma phosphoryl group to a water molecule in the microenvironment [43]. Also, majority of the known kinase inhibitors bind to the ATP binding site, thus abrogating the reaction [44]. An ADP-Glo Kinase assay kit [45] was utilized to determine the kinase activity of the CoV-2 RdRp. The efficiency of the assay in accurately determining a kinase activity was verified using a known kinase – human Akt2. The enzyme exhibited significant activity with a K_m_ value of ∼300 µM for ATP (Fig. 5A), which is very similar to the K_m_ values (∼ 350 µM) determined in previously reported studies [46]. Also, Bovine Serum Albumin, a protein that binds ATP [47] exhibited negligible activity (Fig. 5A), further verifying the specificity and selectivity of the protocol. Notably, incubation of the CoV-2 RdRp with varying concentrations of ATP exhibited significant activity akin to that of the human Akt2, with a K_m_ of ∼500 µM (Fig. 5A). Together, these results present the first ever evidence of an intrinsic kinase/phosphotransferase like activity exhibited by an RdRp molecule from coronaviruses.

**Figure 5.**
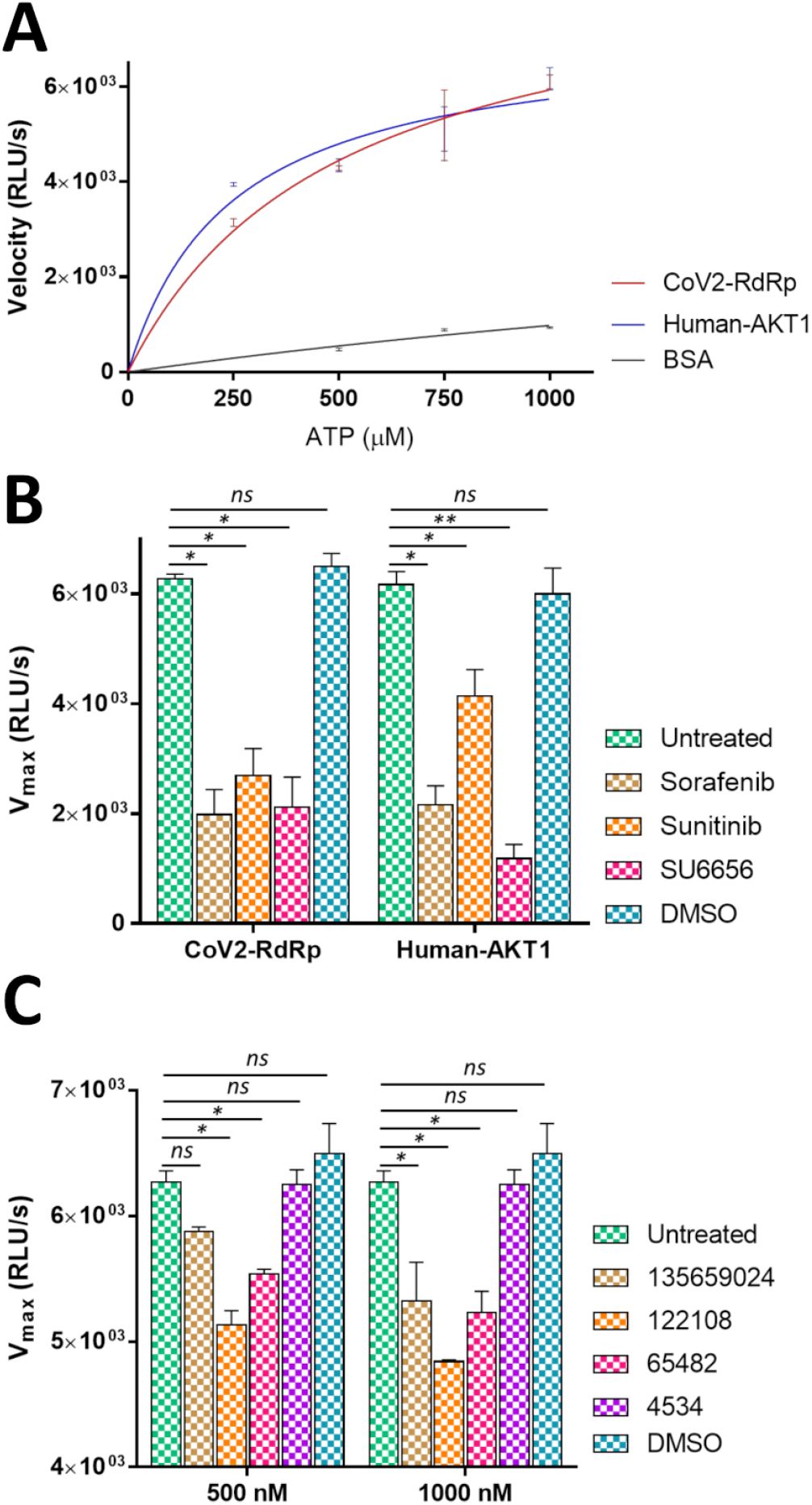
SARS-CoV-2 RdRp exhibits a kinase like activity. **(A)** The CoV-2 RdRp exhibits a kinase like activity akin to that of purified human Akt2. The ATP binding protein Bovine Serum albumin serves as a negative control (Data points show mean and standard error. The connecting curves represent non-linear regressions). **(B)** While treatment with all the kinase inhibitors abrogate the kinase like activity of CoV-2 RdRp, the activity human Akt2 is majorly inhibited by Sorafernib and SU6656 with Sunitinib only exhibiting mild inhibition. **(C)** Treatment of CoV2-RdRp with nucleotidyl transferase inhibitors 135659024, 122108 and 65482 exhibit conspicuous inhibition of the kinase activity in micromolar concentrations. The compound 4534 however fails to exhibit any inhibitory potential (The bars represent mean and standard error. The symbols “^*^” and “^**^” represent significance for p-value less than 0.05 and 0.005, respectively. The symbol “*ns”* represents non-significance).

To further ascertain the possible kinase like activity of CoV-2 RdRp, both CoV-2 RdRp and human Akt2 were incubated with excess of ATP and treated with 500 nM of the each of the three kinase inhibitors mentioned in this study-Sorafenib, Sunitinib and SU6656. Interestingly, all the three kinases inhibitors significantly abrogated the kinase like activity of CoV-2 RdRp (Fig. 5B). For human Akt2, Sorafenib and SU6656 significantly inhibited its kinase activity, while Sunitinib treatment demonstrated a mild inhibition (Fig. 5B). This provides a key insight into drug re-purposing for COVID-19, as all the three aforementioned compounds are well approved drugs for human usage with high tolerance and without any significant toxicity. As the NiRAN domain of CoV-2 RdRp has been annotated as nucleotidyl transferase, we sought to determine any inhibitory efficacy of the top five compounds predicted through *in silico* approaches. We were unable to procure the compound 23673624 commercially, thus we used the four remaining compounds-135659024, 122108, 65482 and 4534. We observed very little effects of the first three compounds on the kinase like activity of CoV-2 RdRp at a concentration of 500 nM (Fig. 5C). However, these compounds which are nucleoside analogs/derivatives exhibited conspicuous inhibitions at a concentration of 1000 nM (Fig. 5C). The fourth compound 4534, a catechol derivative failed to inhibit the CoV-2 RdRp kinase like activity at any of the two concentrations (Fig. 5C). This suggests that the three inhibitory compounds might be involved in blocking the binding of ATP to its respective binding site, in a manner similar to the specific kinase inhibitors.

## Discussion

SARS-CoV-2, the causative agent of the on-going global pandemic COVID-19, is a recently emerged pathogen that has infected over 75 million and caused over 1.6 million deaths as of 20^th^ December, 2020 [48]. With a basic reproduction number ranging from 1.4 to 3.9, the disease quickly spread across the globe within past five months [49]. The situation demands urgent attention from researchers worldwide in order to develop a better understanding of the viral pathogenesis, clinical presentations and biology of the disease in order to develop therapeutics such as vaccines and small molecule inhibitors targeting the viral proteins. Prevalent hypotheses suggest that multiple genome level recombination and zoonotic events between coronaviruses affecting human and bat host have resulted in the evolution of SARS-CoV-2 [15]. However it is worth noticing that the RdRp molecule of SARS-CoV-2 remains largely unchanged at the protein sequence level when compared to previous human coronaviruses such as SARS-CoV, MERS-CoV, SARS-Hku1 as well as non-human coronaviruses like Bat-CoV-Hku4 and Bat-CoV-ZC45, to name a few [50].

Despite being a well-known pathogen, following the SARS-CoV outbreak in 2002, the functional aspects of the RdRp molecule from coronaviruses remains unclear [5,6,11]. In addition, even fewer information is available on the peculiar NiRAN and interface domains that are specific to the RdRp molecules of viruses of the order *Nidovirales* [2,3]. In this study, we propose that the NiRAN domain harbors an atypical kinase/phosphotransferase like fold, with nucleoside triphosphate binding activity. The binding of the wide specificity kinase inhibitors at the predicted active site, further suggest the possible enzymatic action of the NiRAN domain. Interestingly, a recent report has suggested the possibility of kinase inhibitors in management of COVID-19 [51]. In line with the previous study on EAV-RdRp, this study delineates the possible catalytic residues at the NiRAN active site-F35, D36, F48, K73, R116, D218 and D221. In addition, the results also suggest that SARS-CoV-2 RdRp demonstrates a putative kinase/phosphotransferase like activity comparable to that of a well-established human kinase-Akt-1.

Using computational docking and simulation, this study predicts possible inhibitor compounds against the NiRAN domain. Interestingly, the aforementioned kinase like activity of the RdRp is significantly inhibited in the presence of these inhibitors in nanomolar concentrations. The compound Sunitinib is an inhibitor of the tyrosine family kinases [52]; Sorafenib inhibits activities of both serine/threonine and tyrosine family kinases [53,54]; while SU6656 inhibits the Src family kinases [54,55]. While Sorafernib and Sunitinib as approved for medical use in cases of renal, hepatocellular and gastro-intestinal cancers, the compound SU6656 is an experimental molecule used to study the role of Src kinases in cell cycle [56,57]. In addition to the kinase inhibitors, the compound 65482 (Sinefungin), is broad specificity microbial nucleotidyl transferase inhibitor, and is known to inhibit RNA replication in flaviviruses and herpes viruses [58,59]. The compound 122108 (Ribavirin 5’-triphosphate), inhibits the formation of g-capping of RNA in a wide range of viruses, such as Dengue virus, Hantaan virus and Hepatitis C virus [8,60,61]. Officially known as m7GTP, the compound with the PubChem Id 135659024, interferes with the RNA/DNA g-capping activity in many viral pathogens such Rift valley fever virus, influenza virus, Zika virus and dengue virus, to name a few [62–64]. Given the unavailability of SARS-CoV-2 specific therapies and the emergence of newer strains [16], drug repurposing might prove crucial in combating the on-going epidemic while simultaneously being cost and time effective. This study presents well studied drug scaffolds that might be effective against the kinase like catalytic activity of the NiRAN domain of SARS-CoV-2-RdRp. As previous studies have indicated the indispensability of NiRAN catalytic function towards a successful RNA replication in Nidoviruses, it is pertinent to explore *in vivo* anti-viral efficacies of these compounds against coronaviruses.

## Materials and Methods

### Characterization of NiRAN and Interface domains of CoV-2-RdRp

The coordinates of the 3D structure of SARS-CoV-2-RdRp were retrieved from the Protein Data Bank (PDB IDs-7BTF and 6M71) [6]. The dataset 7BTF was used for subsequent analysis as it provided information on all amino acid residues of CoV-2-RdRp. The coordinate data within the file was trimmed to amino acids 1-365 (sequence for NiRAN and Interface domains). This file was used for computational characterization of NiRAN and Interface domains. For sequence analysis, all the coronavirus RdRp sequences were retrieved from the NCBI database and subjected to multiple sequence alignment using the Clustal Omega [65] and Multalin [66] servers. The sequences alignment visualization was generated using the ESPript 3.0 web-tool [67]. Functional analyses were performed using the MPI Bioinformatics Toolkit [17], the DALI server [68] and the DSIMB server [18,69]. The coordinate files for the kinases were retrieved from the Protein Data Bank (PDB IDs-1CSN, 1GAG, 2NRU, 2PVF, 3Q60 and 6EAC) for structural alignment with the NiRAN and Interface domains of CoV-2-RdRp. The alignments were performed using the flexible “jFATCAT” algorithm in the Java application “strucalign” available at the Protein Data Bank [70]. The default alignments parameters were used (RMSD cut off-3 Å, AFP distance cut off-5, Maximum number of twists-5, and Fragment length-8). All visualization were performed in Pymol [71]. Motif prediction and kinase functional prediction were performed using the web servers-ELM [72], PSIPRED [73], CDD [74], MyHits [75], and PhosphoPredict [76].

### *In silico* ligand preparation, active site pocket prediction and molecular docking

The coordinate files for all the inhibitors were retrieved from the PubChem Database and/or from Protein Data Bank. All the inhibitors were prepared by assigning appropriate geometries, bond orders, tautomers and ionization states using LigPrep module implemented in the “Schrodinger Suite 2019” [77]. The coordinate file for the NiRAN and Interface domains was used to predict the probable ligand binding pockets using the SiteMap module of the “Schrodinger Suite 2019”. The accuracy of the sites were verified using the Phyre [78] and CASTp servers [79] and the results were compared. These sites were used to generate a Grid for molecular docking of the inhibitors. The prepared inhibitors were docked onto the predicted binding sites using Glide XP (extra Precision) module of the “Schrodinger Suite 2019”.

### MMGBSA Calculation

The free energies of binding for the inhibitor-domain complexes (NiRAN and Interface domains with inhibitors) were analysed by MMGBSA (Molecular Mechanics Generalized Born Surface Area) calculation [80]. The calculation was executed using the following equation:

G = ΔE_MM_ + ΔG_SGB_ + ΔG_SA_

(ΔE_MM_ represents the minimized molecular energy changes in gas phase; G_SGB_ represents the surface calculation using the GB model; and G_SA_ represents the accessible surface area of the complexes.)

### ADME/T properties prediction

The top 5 inhibitors with the highest docking scores and minimum binding energies were analysed for their pharmacokinetic properties such ADME (Absorption, Distribution, Metabolism, Elimination or Toxicity) and drug likeliness [81]. The QikProp module of the “Schrodinger Suite 2019” was utilized to evaluate the drug likeliness by employing the Lipinski’s rule of five [38] and various pharmacokinetic properties such as probable Hydrogen bonding atoms; Human Oral Absorption (<25 %: poor; >above 80 %: good); QP LogS (Predicted aqueous solubility of the compounds, acceptable range is -6.0 to -0.5) and the IC_50_ values for blockage of the human Ether-a-go-go-Related Gene (hERG) K^+^ channels (< -5: satisfactory)[82].

### DFT Calculation

In quantum mechanics, DFT (density functional theory) calculation is an estimation of the molecular electronic features such as electron density, frontier molecular orbital (HOMO and LUMO) density and electrostatic molecular map in order to predict the reactivity of proposed inhibitors [83,84]. Geometry optimization of compound was performed using hybrid DFT approach at B3LYP (Becke’s three-parameter exchange potential and the Lee–Yang–Parr correlation functional) with 6-31G^**^ basis set [85,86]. The Poisson–Boltzmann solver was utilized to estimate the energy in an aqueous condition that simulates a near physiological state of the target molecule and provides information about the global and local indices of the molecules with respect to their chemical reactivity. All DFT calculations were carried out using Jaguar, version 9.1 [87,88].

### Cloning, Overexpression and Purification of SARS-CoV-2 RdRp

The ORF encoding for the SARS-CoV-2 RdRp was PCR amplified using gene specific primers and using the plasmid pDONR223 SARS-CoV-2 NSP12 as the template DNA (Addgene, USA). Codons encoding for a deca-histidine tag was incorporated into the forward primer in order to obtain the recombinant protein with an N-terminal deca-histidine tag. An entry cloning site (CACC), a recognition site for the restriction endonuclease *NdeI* (CATATG) and a start codon were also incorporated into the forward primer. The reverse primer included a stop codon and the recognition site for the restriction endonuclease *HindIII* (AAGCTT). The primers used are listed at the end of this paragraph. The PCR amplified product was ligated into the entry vector pENTR-D-TOPO using appropriate protocol (Qiagen, USA). The ligated product was transformed into chemically competent *Escherichia coli* strain DH5α and grown on LB agar medium containing 50 µg/ml kanamycin for selection at 310 K. Positive colonies were screened using colony PCR and recombinant plasmid pENTR-D-TOPO-RdRp was extracted by plasmid mini prep using appropriate protocol. The isolated plasmid was subjected to restriction digestion with the enzymes *NdeI* and *HindIII* using appropriate protocol (New England Biolabs, UK). The expression vector pET-28b (+) was linearized by restriction digestion with enzymes *NdeI* and *HindIII*. The digested fragment containing the ORF for RdRp and the linearized expression vector were ligated by T4 DNA ligase employing the appropriate protocol (Qiagen, USA). The ligated product was transformed into chemically competent *Escherichia coli* strain DH5α and grown on LB agar medium containing 50 µg/ml kanamycin for selection at 310 K. Positive colonies were screened using colony PCR and recombinant plasmid pET-28b-RdRp was extracted by plasmid mini prep using appropriate protocol. The plasmids were verified by restriction digestion with enzymes *NdeI* and *HindIII* and by DNA sequencing (BioServe, India). The plasmid pET-28b-RdRp was then into transformed into chemically competent *Escherichia coli* strain BL21-DE3 and grown on LB agar medium containing 50 µg/ml kanamycin for selection at 310 K.

Forward Primer 5’-CACCCATATGAUGCACCATCACCATCACCATCACCACCCAACTTTGTACAAAAAAGTT-3’

Reverse Primer 5’-TATAAGCTTTTACTGCAGCACGGTGTGAGGGGTGTA-3’

A single colony was picked from the agar medium and inoculated into LB liquid medium containing 50 µg/ml kanamycin and grown at 310 K. Large scale liquid cultures were further inoculated from the aforementioned culture and were grown at 310 K. These cultures were induced with 0.5 mM of IPTG at an A_600_ of 0.8. The cultures were maintained for 3 hours post induction at a temperature of 301 K. The cells were harvested by centrifugation of the cultures at 10000 g and were re-suspended in the lysis buffer (20 mM Tris-CL, 300 mM NaCl, 4 mM MgCl_2_, 5 mM 2-mercaptoethanol, 10 % v/v glycerol (Sigma Aldrich, USA) and 0.1 % v/v protease inhibitor cocktail (ThermoScientific, USA) at a pH of 7.0). The cells were lysed using a CF Cell Disruptor (Constant Systems, UK). The lysate was centrifuged at 12000 g for one hour and the supernatant was utilized for protein purification. The protein was purified using the fast performance liquid chromatography system AKTA Pure (GE Healthcare, USA). The protein was purified using an affinity chromatography (HisTrap-FF column; GE Healthcare, USA), followed by an ion exchange chromatography (HiTrap Q HP column; GE Healthcare, USA) and finally by a size exclusion chromatography (HiLoad 16/600 Superdex 200 PG column, GE Healthcare, USA). In affinity chromatography the recombinant protein was eluted in the aforementioned buffer containing 300 mM imidazole (gradient 1 to 500 mM). In ion exchange chromatography the protein eluted in the aforementioned buffer containing an additional 210 mM NaCl (gradient 1 to 1000 mM). In size exclusion chromatography, the protein eluted at a flow volume of ∼65 ml. The complete purification process was maintained at a temperature of 277 K. The purity of the recombinant protein was analysed on SDS-PAGE and the identity of the protein was confirmed using liquid chromatography-tandem mass spectrometry (Ultra-High Resolution Nano LC-MS/MS using C18 column and Q Exactive Orbitrap Mass Spectrometer, ThermoScientific, USA) at a commercial facility (VProteomics, New Delhi, India). The purified protein was concentrated to 1 mg/ml using a 50 kDa membrane centrifugal unit (Millipore, USA) and stored for further use.

### Assessing the kinase/phosphotransferase activity of SARS-CoV-2 RdRp

The ADP-Glo Kinase Assay system (Promega, USA) was utilized for all biochemical analysis using purified recombinant RdRp [45]. The enzyme concentration used for all biochemical assays was 100 nM. The buffer used for biochemical assay contained 20 mM Tris-CL, 300 mM NaCl, 7.5 mM MgCl_2_, 5 mM 2-mercaptoethanol and 10 % v/v glycerol. Varying concentration of ATP (250, 500, 750 and 1000 µM) were utilized to determine the enzymatic activity of the recombinant RdRp. The readouts were measured at 0, 5, 20, 35 and 50 minutes post initiation of the reaction using a luminometer (Berthold Technologies, Germany). Purified Human Akt2 (Sigma Aldrich, USA) at 100 nM and Bovine Serum Albumin at 100 µM (HiMedia, India) concentrations were used as positive and negative controls for Kinase activity, respectively. The kinase inhibitors Sorafenib, Sunitinib and SU6656 (Sigma Aldrich, USA) were dissolved in 100 % v/v DMSO (Sigma Aldrich, USA) and used at a concentration of 500 nM. The predicted nucleotidyl transferase inhibitors 135659024, 122108, 65482 and 4534 (Sigma Aldrich, USA) were dissolved in 100 % v/v DMSO and used at two different concentrations-500 and 1000 nM. For all inhibition studies, the ATP concentration was maintained at 100 mM. The graphs were plotted using GraphPad Prism 7.0. Non-linear regressions of the enzymatic activities were performed for obtaining the Michaelis-Menten kinetics. Statistical significance for the inhibitory potential of the compounds was determined using student’s t-test.

## Acknowledgements

This work was supported by the core funding from the National Institute of Immunology, New Delhi, India. Dr. J. Jeyakanthan gratefully acknowledges the MHRD-RUSA 2.0 [F.24/51/2014-U, Policy (TNMulti-Gen), Dept. of Edn., Govt. of India] for the infrastructure facilities provided to the Department of Bioinformatics, Alagappa University.

## Author Contributions

AD, RM, MA, SC, DK and ST contributed equally to the manuscript. AD and BKB conceptualized the study. AD, RM and MA performed the computational analysis. AD, SC, DK and ST purified the protein and performed the biochemical experiments. AD and BKB analysed the data and drafted the manuscript. All authors proofread and approved the manuscript.

