## Supplemental Figures and Tables for "Characterization of the NiRAN domain from RNA-dependent RNA polymerase provides insights into a potential therapeutic target against SARS-CoV-2"

### Supplementary Figures

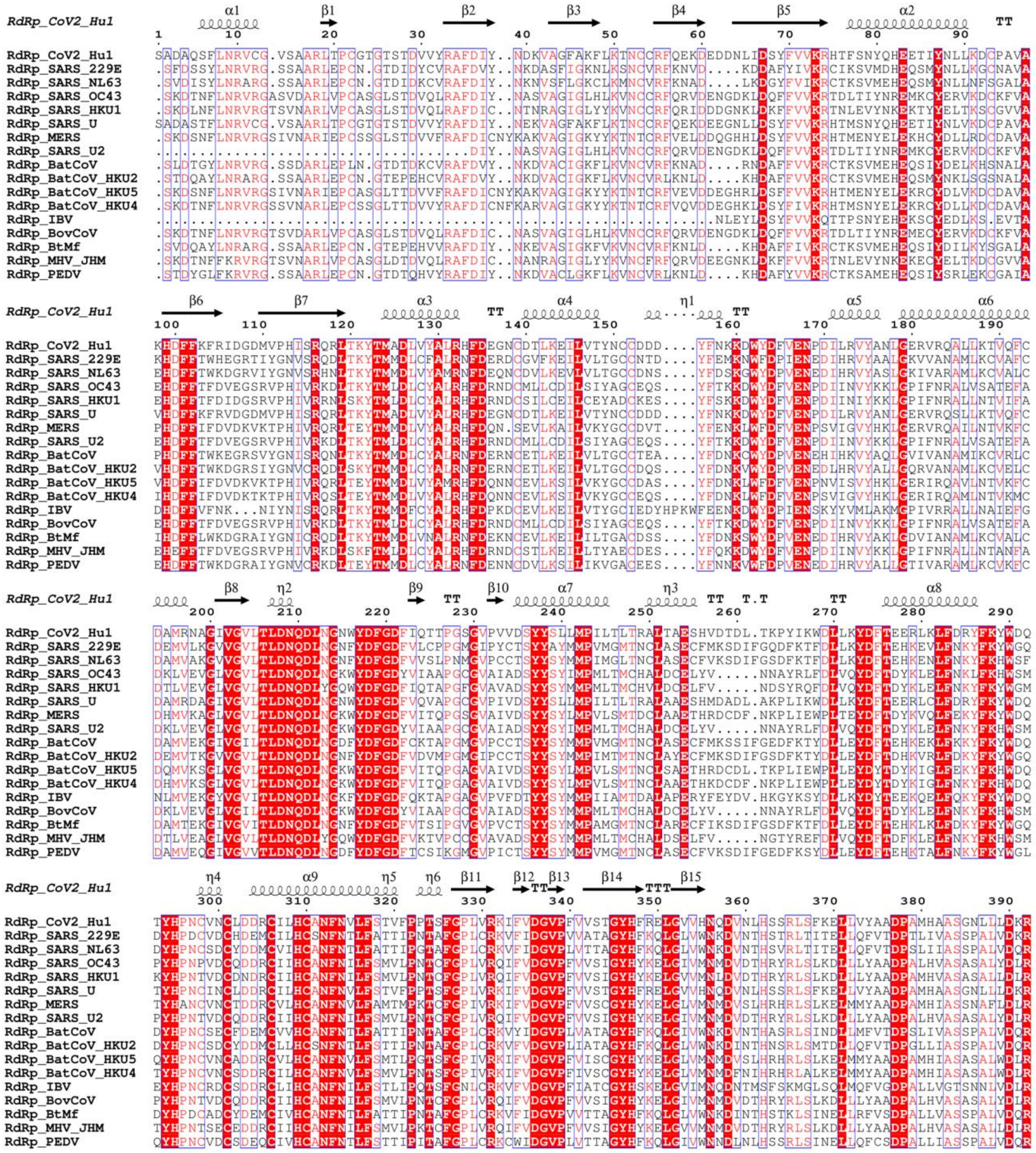

**Figure S1: A multiple protein sequence alignment of NiRAN and interface domains from the RdRp molecules of coronaviruses affect various vertebrate hosts.**

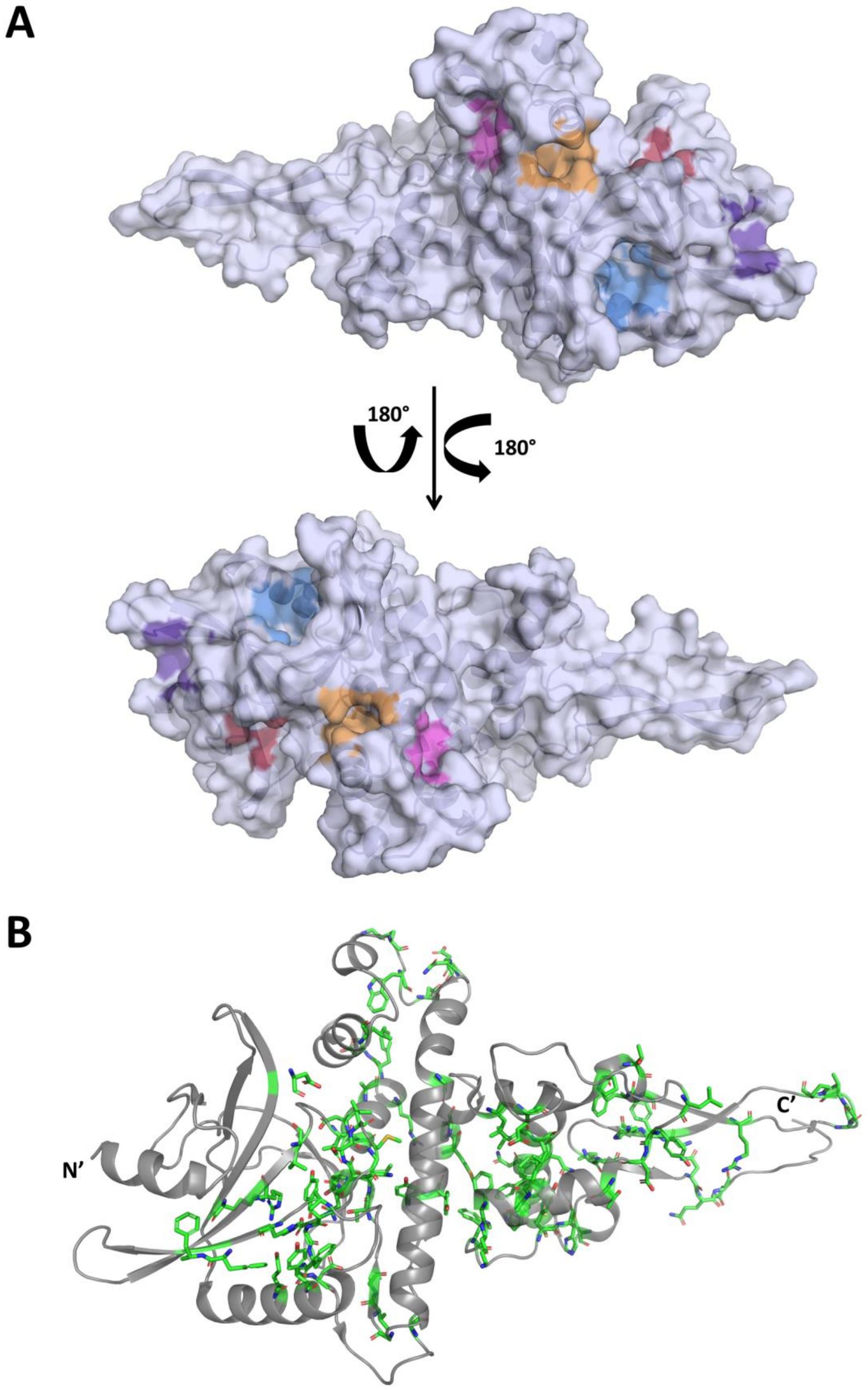


**Figure S2: Organization and conservation of the NiRAN Domain. (A)** Organization of the entry pockets at CoV-2-RdRp NiRAN domain lined with the strictly conserved residues**. (B)** While the conserved residues in the interface domain are scattered across the structural elements, the majority of the conserved residues in NiRAN domain lie between the antiparallel β-sheet and the immediately following helix bundle, possibly hinting at the NiRAN domain active site.

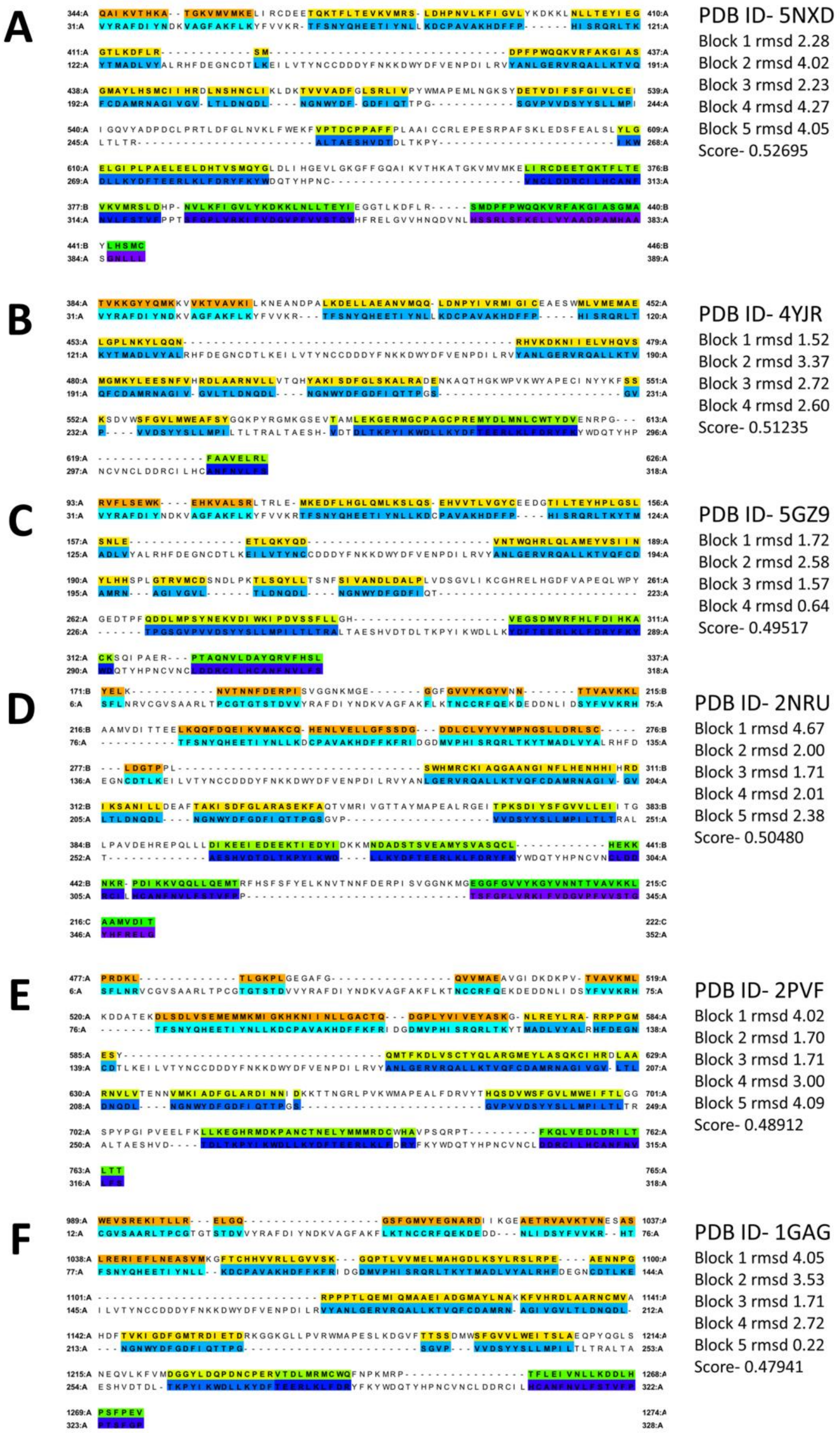


**Figure 3 (Legend on next page)**

**Figure S3: Pairwise alignments corresponding to the alignment of the structural elements of CoV-2-RdRp NiRAN with known Kinases. (A)** Lim 2 kinase domain. **(B)** Syk kinase domain **(C)** O-mannosyl kinase domain. **(D)** IRAK4 kinase domain. **(E)** FGFR2 kinase domain. **(F)** Insulin receptor kinase domain. (Colour codes- Block 1: orange-cyan; Block 2: yellow-turquoise; Block 3: lime- deep blue; Block 4: green- deep blue; Block 5: green- purple)

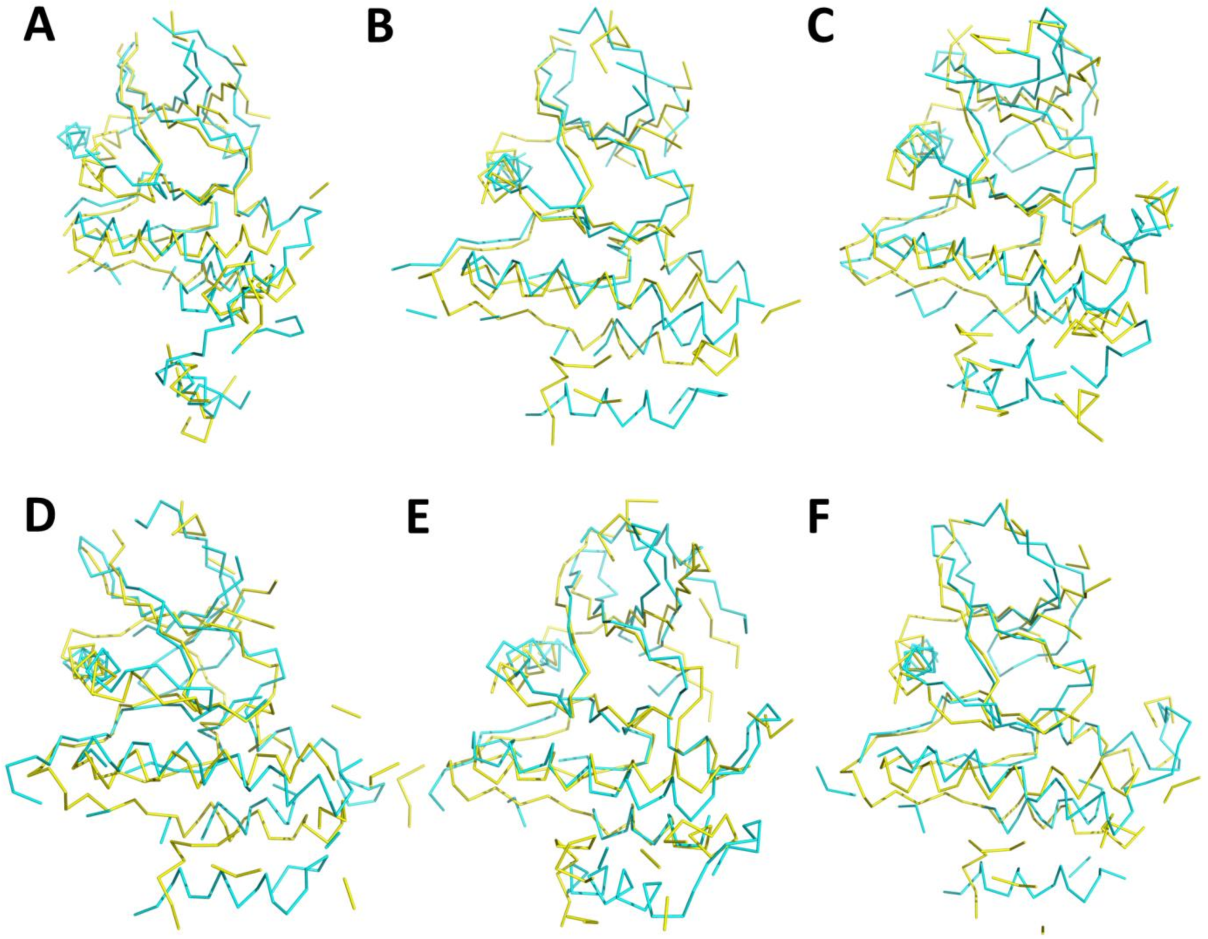


**Figure S4: The structural superimposition of the Cα chains of CoV-2-RdRp NiRAN with known Kinases reveal significant alignment of the polypeptide chains. (A)** Lim 2 kinase domain. **(B)** Syk kinase domain **(C)** O-mannosyl kinase domain. **(D)** IRAK4 kinase domain. **(E)** FGFR2 kinase domain. **(F)** Insulin receptor kinase domain. (The aforementioned kinases' Cα chains are shown in cyan, CoV-2-RdRp NiRAN Cα chains are shown in yellow.)

>RdRp_CoV2_NiRAN_Interface

SADAQSFLNRVCGVSAARLTPCGTGTSTDVVYRAFDIYNDKVAGFAKFLKTNCCRFQE**KDEDDNLIDSYFVVKR**

**HTFSNYQHEETIYNLLKDCPAVAKHDFFKFRIDGDMVPHISRQRLTKYTMADLVYALRHFDEGNCDTLKEILVTY**

**NCCDDDYFNKKDWYDFVENPDILRVYANLGERVRQALLKTVQFCDAMRNAGIVGVLTLDNQDLNGNWYDFGD**

**FIQTTPGSGVPVVDSYYSLLMPILTLTR***ALTAESHVDTDLTKPYIKWDLLKYDFTEERLKLFDRYFKYWDQTYHPN*

*CVNCLDDRCILHCANFNVLFSTVFPPTSFGPLVRKIFVDGVPFVVSTGYHFRELGVVHNQDVNLHSSR*

**Figure S5: The prediction of kinase like motifs reveals the presence of sequence motifs belonging to kinase families PkA, PkC and Src.** A kinase consensus site search prediction also predicts multiple phosphorylation sites. (Bold- NiRAN domain; Italics- Interface domain; Underlines: Red- Kinase consensus sequence, Blue- PkC like motif, purple- PkA like motif, Green- Src kinase like motif, Grey- Unspecified Kinase like motifs, Yellow- Myristoylation site consensus sequence.)

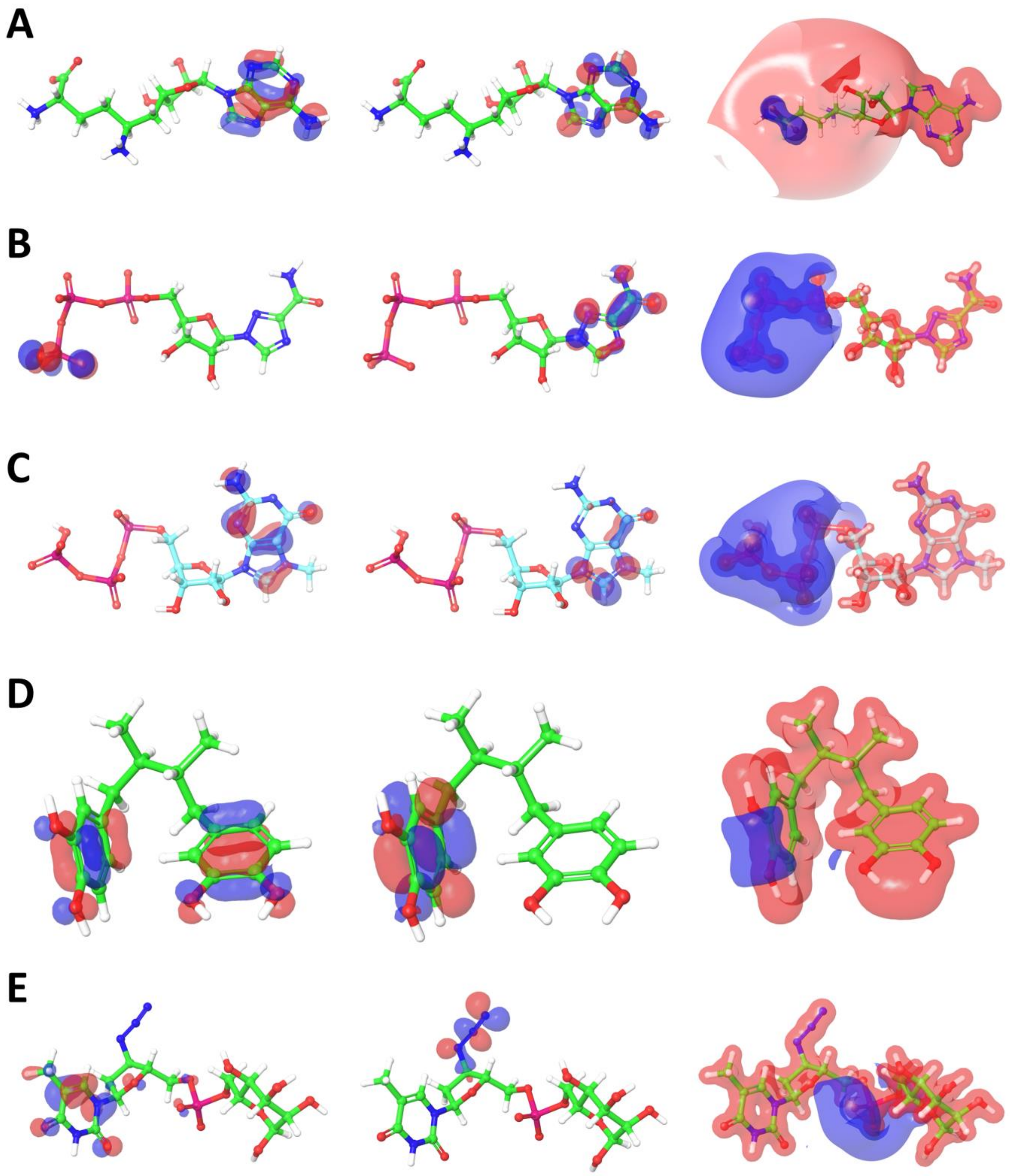


**Figure S6: The DFT predictions of the best five predicted nucleotidyl transferase inhibitors. (A)** 65482. **(B)** 122108. **(C)** 135659024. **(D)** 4534. **(E)** 23673624. (Left panels represent HOMO; Centre panels represent LUMO; and Right panels represent Electrostatic potential surface density.)

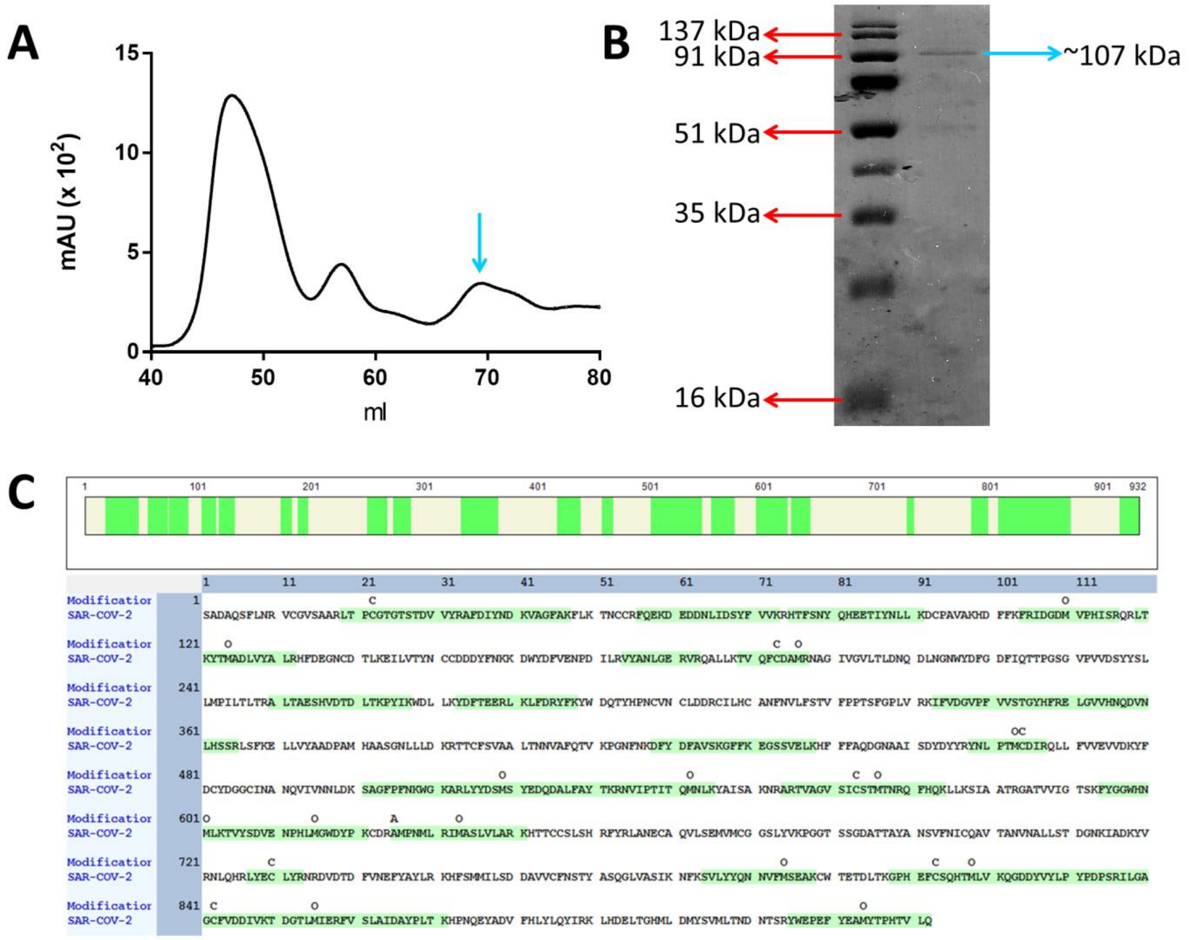


**Figure S7: Purification of CoV2-RdRp. (A)** Size exclusion chromatogram of recombinant SARS-CoV-2 RdRp (The peak indicated with a blue arrow represents the purified protein sample). **(B)** SDS-PAGE profile of the purified SARS-CoV-2 RdRp (The band indicated with a blue arrow represents the purified protein sample and the molecular weight markers are indicated in red arrows).

#### Supplementary Tables

**Table S1: Top 15 molecules with structurally similar fold(s) predicted by HHPred analysis.**

| No. | Hit (PDB) | Prob | E-value | P-value | Score | SS | Query HMM | Template HMM |
|---|---|---|---|---|---|---|---|---|
| 1 | 6NUR | 100.0 | 4.2E-72 | 8.4E-77 | 552.9 | 19.6 | 1-190 | 62-251 |
| 2 | 7BTF | 100.0 | 6E-69 | 1.2E-73 | 528.8 | 20.6 | 1-190 | 60-249 |
| 3 | 4YJR | 86.2 | 5.7 | 0.00011 | 32.1 | 7.9 | 4-161 | 38-160 |
| 4 | 2H34 | 86.2 | 7.4 | 0.00015 | 32.8 | 8.9 | 8-161 | 59-179 |
| 5 | 6INY | 82.7 | 2.6 | 5.1E-05 | 40.0 | 5.1 | 117-161 | 217-262 |
| 6 | 6EAC | 82.7 | 2.2 | 4.3E-05 | 40.6 | 4.7 | 117-161 | 220-265 |
| 7 | 5GZ9 | 81.4 | 29 | 0.00057 | 29.5 | 10.6 | 8-172 | 57-195 |
| 8 | 5DRB | 81.1 | 11 | 0.00021 | 30.8 | 7.6 | 8-161 | 52-178 |
| 9 | 6BXI | 81.1 | 13 | 0.00026 | 32.1 | 8.5 | 8-161 | 31-151 |
| 10 | 5NXD | 79.1 | 18 | 0.00036 | 30.3 | 8.6 | 9-161 | 28-144 |
| 11 | 6TU9 | 77.8 | 32 | 0.00063 | 29.0 | 9.7 | 4-161 | 44-183 |
| 12 | 6FEX | 76.9 | 36 | 0.00072 | 28.8 | 9.8 | 8-161 | 57-188 |
| 13 | 3M2W | 74.7 | 30 | 0.00059 | 28.7 | 8.6 | 8-161 | 43-165 |
| 14 | 6CQE | 74.5 | 42 | 0.00083 | 27.4 | 9.3 | 9-161 | 42-158 |
| 15 | 4BTF | 74.5 | 24 | 0.00048 | 31.2 | 8.4 | 4-161 | 214-343 |

**Table S2: Top 15 molecules with structurally similar fold(s) predicted by ORION_DSIMB analysis.**

| No. | Score | Ungap_score | Pval_Query | Pval_Target | Query_Start_End | Target_Start-end | HIT (PDB) |
|---|---|---|---|---|---|---|---|
| 1 | 57.323 | 43.106 | 1.64E-09 | 1.76E-112 | 1-181 | 1-192 | 3H93 |
| 2 | 56.250 | 47.533 | 7.00E-09 | 5.15E-112 | 1-189 | 1-214 | 3BWY |
| 3 | 55.739 | 46.048 | 1.35E-08 | 8.59E-112 | 3-190 | 1-223 | 1G7S |
| 4 | 55.713 | 49.929 | 1.39E-08 | 8.82E-112 | 6-188 | 1-185 | 1GBS |
| 5 | 54.632 | 48.730 | 5.18E-08 | 2.60E-111 | 15-188 | 1-181 | 3MGW |
| 6 | 54.567 | 47.204 | 5.59E-08 | 2.77E-111 | 1-186 | 1-189 | 1U6M |
| 7 | 53.917 | 44.721 | 1.17E-07 | 5.31E-111 | 18-173 | 1-154 | 1DCQ |
| 8 | 53.779 | 41.717 | 1.37E-07 | 6.10E-111 | 1-190 | 1-209 | 2FR1 |
| 9 | 53.320 | 44.212 | 2.26E-07 | 9.65E-111 | 1-188 | 1-198 | 3C3P |
| 10 | 53.242 | 47.062 | 2.46E-07 | 1.04E-110 | 14-189 | 1-177 | 1EB6 |
| 11 | 53.233 | 44.408 | 2.48E-07 | 1.05E-110 | 2-190 | 1-247 | 2YVL |
| 12 | 53.060 | 45.677 | 2.98E-07 | 1.25E-110 | 21-189 | 1-157 | 3TWR |
| 13 | 52.739 | 43.585 | 4.17E-07 | 1.73E-110 | 1-190 | 1-225 | 3C3Y |
| 14 | 52.364 | 45.722 | 6.12E-07 | 2.51E-110 | 31-187 | 1-187 | 2CVB |
| 15 | 52.343 | 44.148 | 6.25E-07 | 2.56E-110 | 1-190 | 1-230 | 1G8S |

**Table S3: The complete list of compounds selected for docking at the predicted active site of CoV-2-RdRp NiRAN domain.**

| | NAME | PUBCHEM ID | PDB ID(s) | STRUCTURE 2D |
|---|---|---|---|---|
| **1** | Nootkatin | 238797 | | 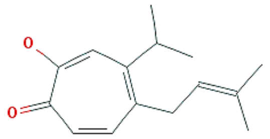 |
| **2** | NSC 282885 | 323364 | | 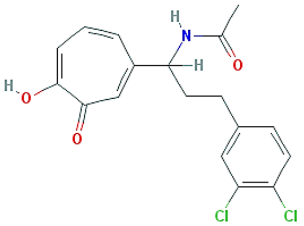 |
| **3** | Manicol | 5351306 | 3QLH | 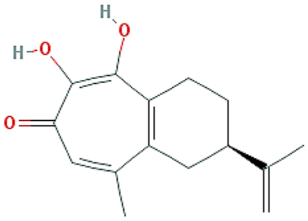 |
| **4** | 2,4,6-Trimethyl-7-oxocyclohepta-1,3,5-trien-1-YL 4-chlorobenzoate | 873418 | | 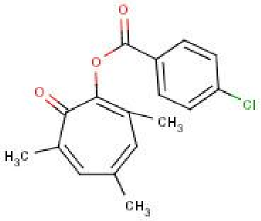 |
| **5** | 7-Oxocyclohepta-1,3,5-trien-1-yl 4-methylbenzenesulfonate | 562886 | | 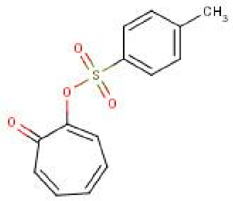 |
| **6** | NSC 79555 | 254856 | | 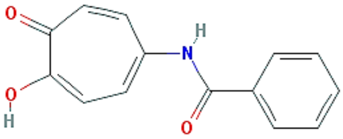 |
| **7** | Labotest 72543251 | 14027158 | | 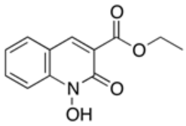 |
| **8** | N-Hydroxy-1,8-naphthalimide | 82263 | | 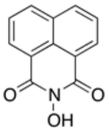 |

| | | | | |
|---|---|---|---|---|
| **9** | (9-(4-Ethoxyphenyl)-2,6,7-trihydroxy-xanthen-3-one) | 2931798 | | 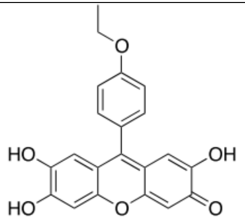 |
| **10** | 9-(4'-Dimethylaminophenyl)-2,6,7-trihydroxyfluorone Sulfate Hydrate | 90475469 | | 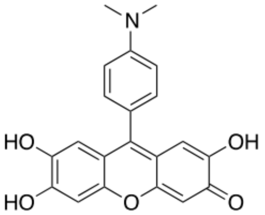 |
| **11** | 9-METHYL-2,3,7-TRIHYDROXY-6-FLUORONE | 72721 | | 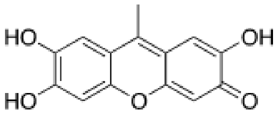 |
| **12** | Asinex BAS0223612 | | | 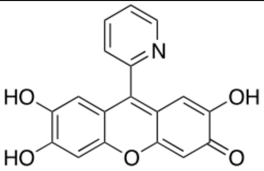 |
| **13** | Elvitegravir | 5277135 | 3L2U, 3L2W | 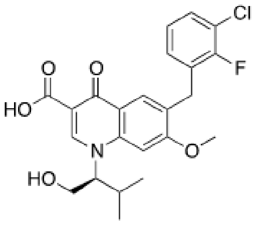 |
| **14** | Raltegravir | 54671008 | 4MDA, 4MDB, 3L2V, 3OYA | 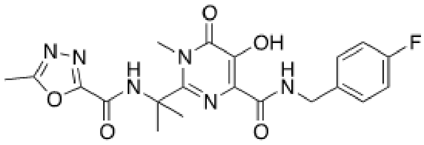 |
| **15** | 7-Chloro-1-cyclopropyl-6-fluoro-1,4-dihydro-4-oxoquinoline-3-carboxylic acid | 483180 | | 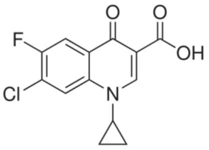 |
| **16** | Ethyl 5-({[(5-benzyl-1,3,4-oxadiazol-2-yl)thio]acetyl}amino)-4-cyano-3-methyl-2-thiophenecarboxylate | 2222655 | | 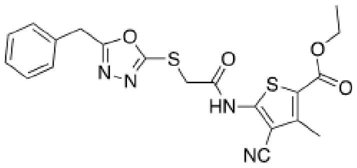 |
| **17** | Ethyl 4-cyano-3-methyl-5-{[(4H-1,2,4-triazol-3-ylthio)acetyl]amino}-2-thiophenecarboxylate | 2193853 | | 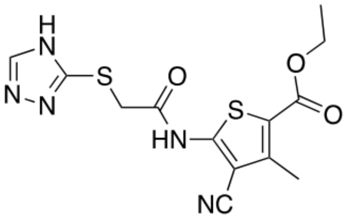 |

| | | | | |
|---|---|---|---|---|
| **18** | Ethyl 5-[[2-[[5-(4-chlorophenyl)-1,3,4-oxadiazol-2-yl]sulfanyl]acetyl]amino]-4-cyano-3-methylthiophene-2-carboxylate | 2224333 | | 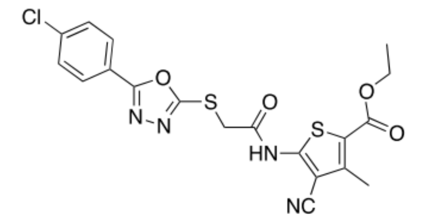 |
| **19** | Ethyl 4-cyano-3-methyl-5-({[(4-methyl-2-pyrimidinyl)thio]acetyl}amino)-2-thiophenecarboxylate | 2198264 | | 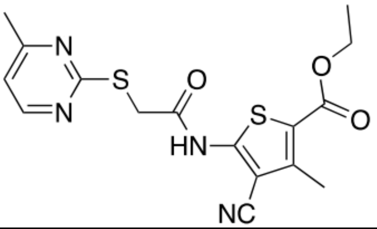 |
| **20** | Nordihydroguaiaretic acid | 4534 | 4PWJ | 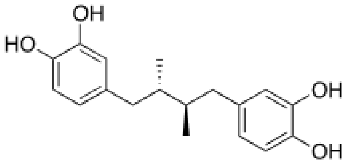 |
| **21** | 5,6-Dimethyl-2-(4-nitrophenyl)thieno[2,3-d]pyrimidin-4(3H)-one | 5345286 | | 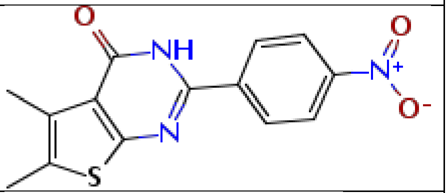 |
| **22** | Acyclovir | 135398513 | 3KD1, 3KD5, 1VZV, 1PWY, 5I3C | 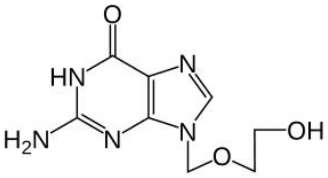 |
| **23** | Ribavirin 5'-triphosphate | 122108 | 1R6A, 2E9R | 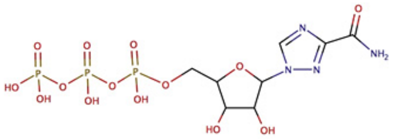 |
| **24** | (Z)-3-Amino-5-(4-chlorobenzylidene)-2-thioxothiazolidin-4-one | 1379483 | | 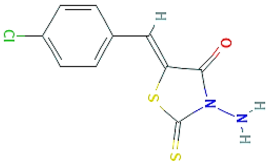 |
| **25** | MADTP | 23673624 | | 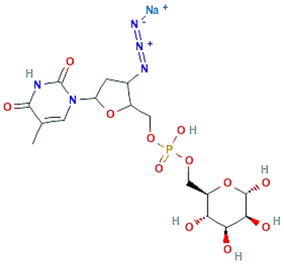 |
| **26** | Sinefungin | 65482 | 5YNB, 5YNN, 5YNP, 5MRK, 4R8S | 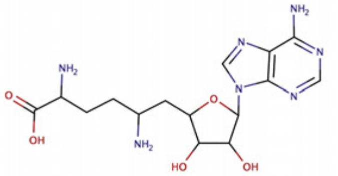 |

| | | | | |
|---|---|---|---|---|
| **27** | NSC 293161 | 325174 | | 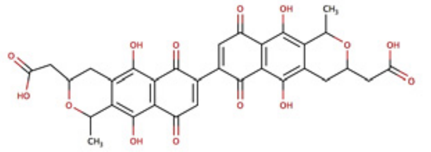 |
| **28** | NSC 23217 | 3246089 | | 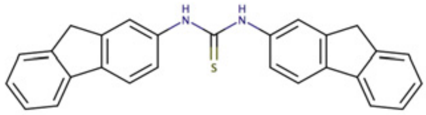 |
| **29** | F3043-0013 | 16644047 | | 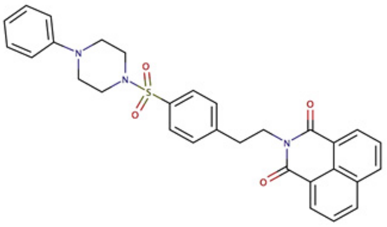 |
| **30** | NSC 125910 | 277277 | | 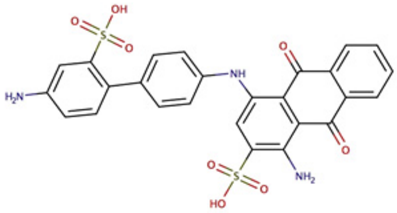 |
| **31** | NSC 371880 | 340799 | | 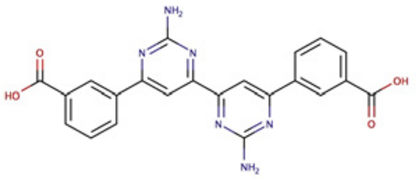 |
| **32** | NSC 372295 | 3246679 | | 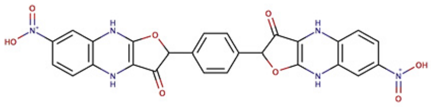 |
| **33** | F0922-0796 | 16193614 | | 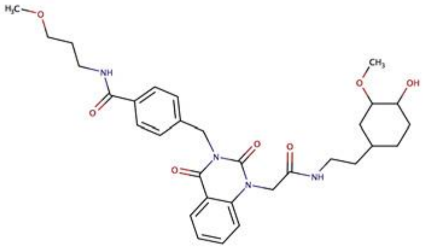 |
| **34** | 3-Deazaneplanocin | 73087 | | 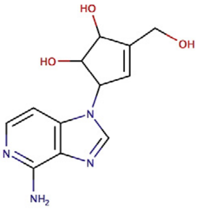 |
| **35** | EGCG | 65064 | 4AWM | 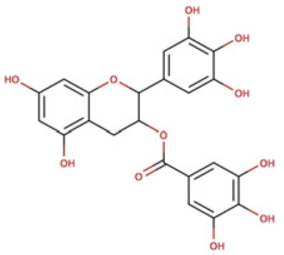 |

| | | | | |
|---|---|---|---|---|
| **36** | (5Z)-2-[2-(2-oxoindol-3-yl)hydrazinyl]-5-(2-oxo-1H-indol-3-ylidene)-1,3-thiazol-4-one | 1554535 | | 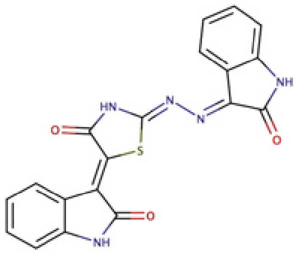 |
| **37** | Flutimide | 6443207 | 1WL8 | 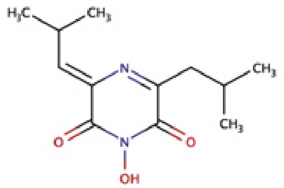 |
| **38** | m7GTP | 135659024 | 6QHG, 6EVK, 5FMM, 5EFA, 4NCE, 4EQK | 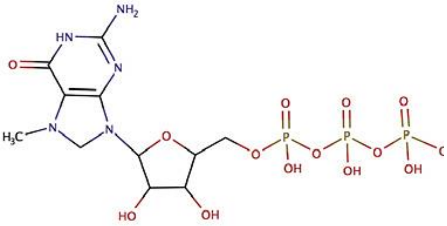 |
| **39** | VX-787 | 126970300 | 5WL0, 6EUV, 4P1U | 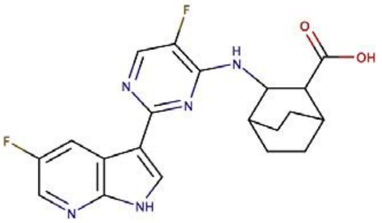 |
| **40** | [4-(Aminomethyl)phenyl]methanesulfonamide | 9167228 | | 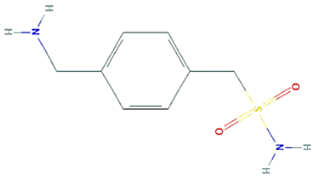 |
| **41** | 9-(4-Hydroxybutyl)-N2-phenylguanine | 135415618 | | 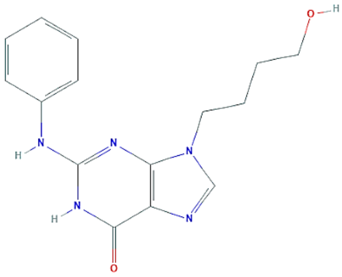 |
| **42** | 1-Hydroxy-2(1H)-quinolinone | 264295 | | 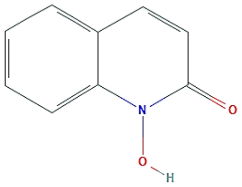 |
| **43** | 2,4,6-Trimethyl-7-oxocyclohepta-1,3,5-trien-1-YL 4-chlorobenzoate | | | 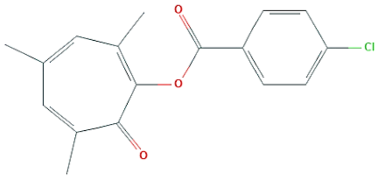 |

| | | | | |
|---|---|---|---|---|
| **44** | 2-Hydroxyisoquinoline-1,3(2H,4H)-dione | 514100 | | 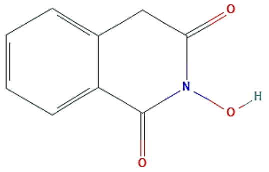 |
| **45** | Ciclopirox | 2749 | 6J10 | 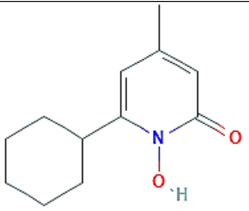 |
| **46** | 7-Oxocyclohepta-1,3,5-trien-1-YL benzoate | 767419 | | 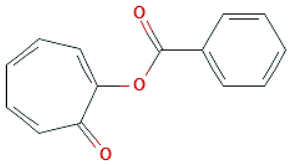 |
| **47** | beta-Thujaplicinol | 72605 | 3IG1, 3V1R, 5C2F | 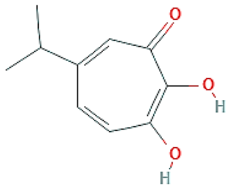 |
| **48** | 1,6-Dibenzyl-5-ethyl-3-hydroxy-pyrimidine-2,4-dione | 52936423 | | 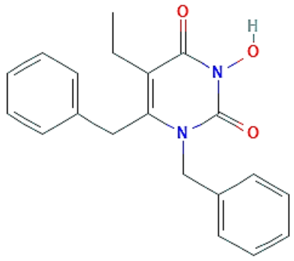 |
| **49** | 6-(3,5-Dimethylbenzoyl)-1-(ethoxymethyl)-5-ethyl-3-hydroxy-pyrimidine-2,4-dione | 52918273 | | 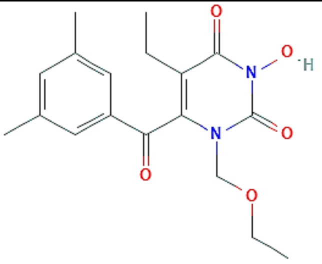 |
| **50** | 5-Benzyl-6-ethyl-1-[(4-fluorophenyl)methoxymethyl]-3-hydroxy-pyrimidine-2,4-dione | 52936326 | | 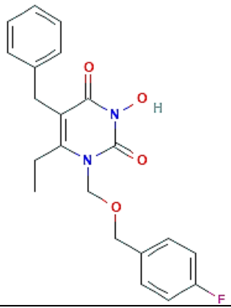 |

| | | | | |
|---|---|---|---|---|
| **51** | 6-Benzyl-5-ethyl-1-[(4-fluorophenyl)methyl]-3-hydroxy-pyrimidine-2,4-dione | 52936424 | | 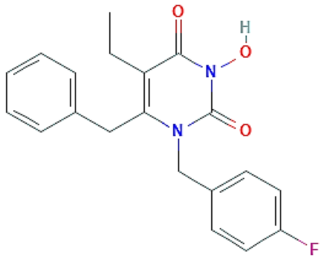 |
| **52** | 6-(3,5-Dimethylbenzoyl)-5-ethyl-1-[2-(4-fluorophenyl)ethoxymethyl]-3-hydroxy-pyrimidine-2,4-dione | 54587147 | | 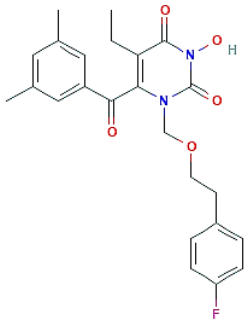 |
| **53** | 1-(Ethylamino)-3-hydroxypyrimidine-2,4-dione | 68725888 | | 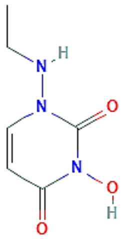 |
| **54** | 5-Benzyl-1-(ethoxymethyl)-6-ethyl-3-hydroxypyrimidine-2,4-dione | 56959484 | | 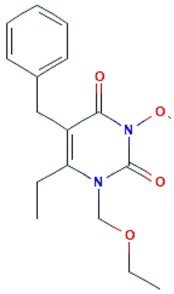 |
| **55** | 6-(3,5-Dimethylphenoxy)-1-(ethoxymethyl)-5-ethyl-3-hydroxypyrimidine-2,4-dione | 56959332 | | 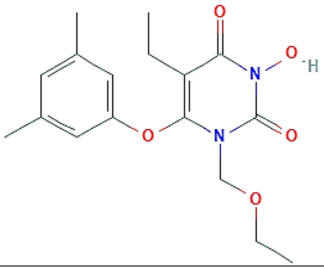 |
| **56** | 6-Ethyl-1-[(4-fluorophenyl)methoxymethyl]-5-[(4-fluorophenyl)methyl]-3-hydroxypyrimidine-2,4-dione | 56959779 | | 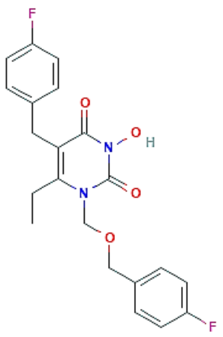 |

| | | | | |
|---|---|---|---|---|
| **57** | 6-(3,5-Dimethylphenoxy)-5-ethyl-1-[(4-fluorophenyl)methoxymethyl]-3-hydroxypyrimidine-2,4-dione | 56959481 | | 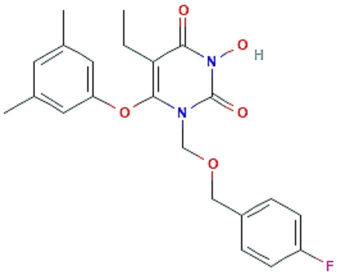 |
| **58** | 1-[(2R,3R,4S,5R)-3,4-Dihydroxy-5-(hydroxymethyl)oxolan-2-yl]-3-hydroxypyrimidine-2,4-dione | 14330990 | | 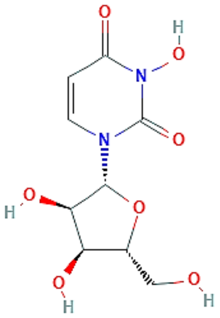 |
| **59** | 6-(Cyclohexylmethyl)-1-[(3,4-difluorophenyl)methoxymethyl]-5-ethyl-3-hydroxypyrimidine-2,4-dione | 129908680 | | 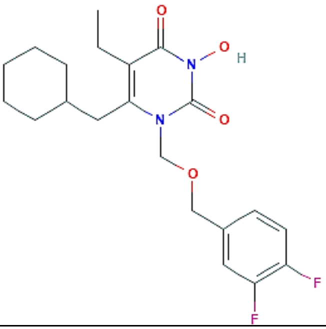 |
| **60** | 1-[(3-Chloro-2-fluorophenyl)methoxymethyl]-6-(cyclohexylmethyl)-5-ethyl-3-hydroxypyrimidine-2,4-dione | 129908679 | | 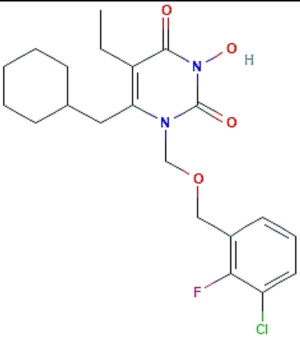 |
| **61** | 1-[(2S,5S)-3-Hydroperoxy-4-hydroxy-5-(hydroxymethyl)oxolan-2-yl]-3-hydroxypyrimidine-2,4-dione | 90111589 | | 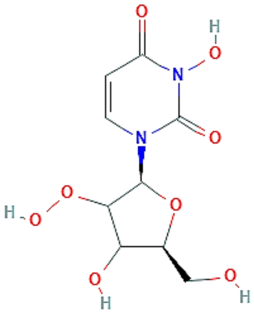 |
| **62** | Sun B8155 | 135484493 | | 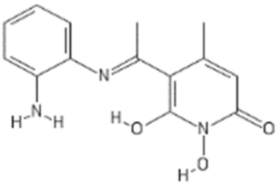 |

| | | | | |
|---|---|---|---|---|
| **63** | Rilopirox | 71778 | 6J10 | 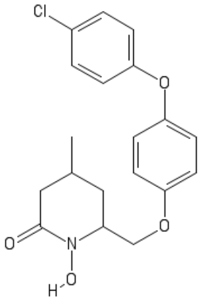 |
| **64** | N-[(4-Fluorophenyl)methyl]-2-hydroxy-7-methoxy-1,3-dioxo-4H-isoquinoline-4-carboxamide | 57389495 | | 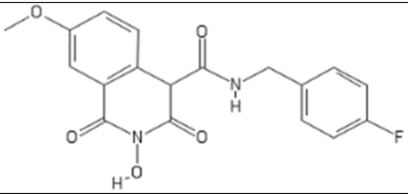 |
| **65** | 7-Benzamido-N-[(4-fluorophenyl)methyl]-2-hydroxy-1,3-dioxo-4H-isoquinoline-4-carboxamide | 57389416 | | 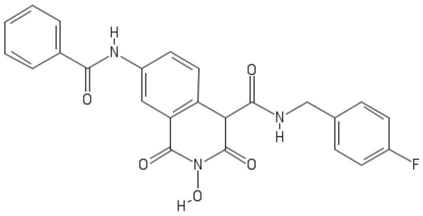 |
| **66** | 2,3-dihydroxy-N-[(4-methoxyphenyl)methyl]-7-nitro-1-oxo-isoquinoline-4-carboxamide | 163671823 | | 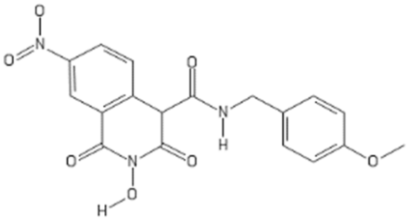 |
| **67** | 7-Chloro-N-[(4-fluorophenyl)methyl]-2-hydroxy-1,3-dioxo-4H-isoquinoline-4-carboxamide | 67984418 | | 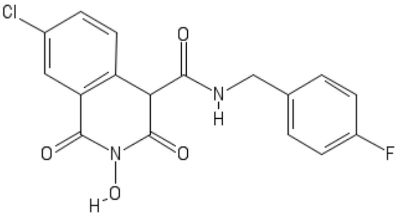 |
| **68** | Ethyl 1-hydroxy-2-oxo-1,2-dihydroquinoline-3-carboxylate | 14027158 | | 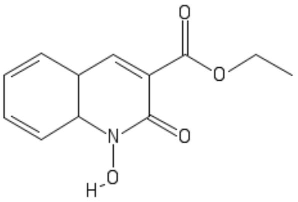 |
| **69** | (5Z)-2-Hydroxy-4-methyl-6-oxo-5-[(5-phenylfuran-2-yl)methylidene]-5,6-dihydropyridine-3-carbonitrile | 2275754 | | 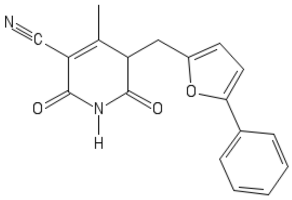 |
| **70** | N-Hydroxyphthalimide | 10665 | | 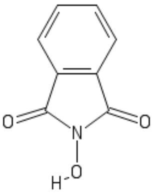 |

| | | | | |
|---|---|---|---|---|
| **71** | 2,3-dihydroxyquinoxaline | 27491 | 5U8C (subset) | 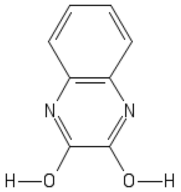 |
| **72** | 3-(alpha-Hydroxy-4-nitrobenzyl)-2,5-dihydroxy-2,4,6-cycloheptatrien-1-one | 13924966 | | 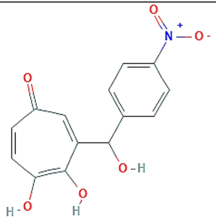 |
| **73** | 3-(alpha-Hydroxy-4-nitrobenzyl)-2,5-dihydroxy-2,4,6-cycloheptatrien-1-one | 13924966 | | 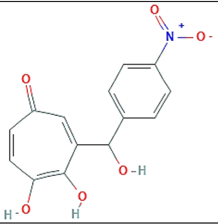 |
| **74** | 1-Hydroxy-4H-pyridine-2,3-dione | 139594038 | | 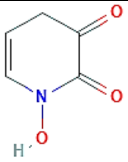 |
| **75** | 1,5-Naphthyridin-2(1H)-one | 589680 | 5TVT, 4BW4 (subset) | 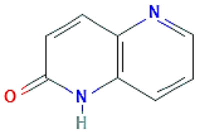 |
| **76** | N-Hydroxy-2-[4-[4-(2-hydroxyethylamino)phenyl] phenyl]-N-methyl-1,6-naphthyridine-4-carboxamide | 77106704 | | 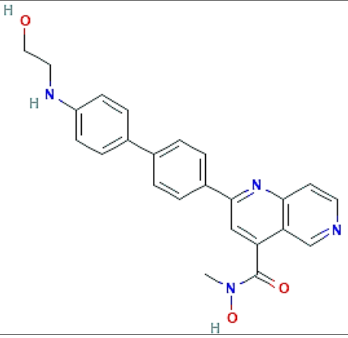 |
| **77** | 5-Bromo-N-(4-fluorobenzyl)-8-hydroxy-1,6-naphthyridine-7-carboxamide | 22346401 | | 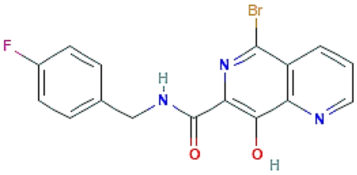 |

**Table S4: Docking analysis of kinase inhibitors at the proposed active site of NiRAN domain.**

| No. | Name | PubChem ID | Docking Score (Kcal/Mol) | Binding Energy (ΔG bind) | H-bond Interactions |
|---|---|---|---|---|---|
| **1** | Sunitinib | 5329102 | -3.50 | -28.07 | Asp36, Asp218 and Asp221 |
| **2** | Sorafenib | 216239 | -3.06 | -36.06 | Asp36 |
| **3** | SU6656 | 5312137 | -2.61 | -25.07 | Lys73 |

**Table S5: Docking analysis of best 5 proposed inhibitors at the proposed active site of NiRAN domain.**

| **No.** | **PubChem ID** | **Docking Score (Kcal/Mol)** | **Binding Energy (ΔG bind)** | **H-bond Interactions** | **Other Interactions** |
|---|---|---|---|---|---|
| 1 | 65482 | -10.35 | -25.38 | Asp36, Asn52, Lys73, Asp218, Asp221 | Phe35 (Pi-Pi), Phe48 (Pi-Pi), Glu83 (Salt Bridge) |
| **2** | 122108 | -7.42 | -10.03 | Lys73, Asp36, Asn52, Asp218, Asp221 | Lys73 (Salt Bridge) |
| **3** | 135659024 | -6.94 | -24.33 | Asn79, Thr76, Arg74, Asn52, Val204, Asn209, | Lys73 (Salt bridge), Asp218 (Salt bridge) |
| **4** | 4534 | -6.84 | -32.07 | Lys73, Asn52, Asp218 | NA |
| **5** | 23673624 | -6.76 | -34.18 | Arg33, Thr51, Asn52, | Lys73 (Salt bridge), Arg116 (Salt Bridge) |

**Table S6: The predicted ADME/T properties of best 5 proposed inhibitors.**

| No. | PubChem Id | Mol.Wt. | H-bond donor | H-bond acceptor | QP LogS | QPLog HERG | Lipinski rule of 5 |
|---|---|---|---|---|---|---|---|
| 1 | 65482 | 381.3 | 9 | 13 | -0.11 | -4.52 | 2 |
| 2 | 122108 | 484.1 | 4 | 21 | -0.00 | 2.26 | 2 |
| 3 | *135659024 | 537.2 | - | - | - | - | - |
| 4 | 4534 | 302.3 | 4 | 3 | -3.39 | -4.55 | 0 |
| 5 | 23673624 | 509.3 | 6 | 21 | -1.62 | -2.95 | 3 |

* The program failed to predict the ADME/T properties

**Table S7: The DFT predictions for the best 5 proposed inhibitors.**

| No. | Compounds ID | HOMO (eV) | LUMO (eV) | $E_{HOMO}$-$E_{LUMO}$ (eV) |
|---|---|---|---|---|
| 1 | 65482 | -0.228 | -0.034 | -0.194 |
| 2 | 122108 | -0.196 | -0.051 | -0.145 |
| 3 | 135659024 | -0.211 | -0.047 | -0.164 |
| 4 | 4534 | -0.208 | -0.002 | -0.206 |
| 5 | 23673624 | -0.243 | -0.044 | -0.199 |

**Table S8: LC-MS/MS verification of the SARS-CoV-2 RdRp.**

| | |
|---|---|
| **Accession** | 6YYT_A |
| **Description** | Chain A, nsp12, Hillen, H.S., *et al.* |
| **Coverage [%]** | 45 |
| **Contaminant** | FALSE |
| **Peptides** | 41 |
| **PSMs** | 83 |
| **Unique Peptides** | 41 |
| **Protein Groups** | 1 |
| **AAs** | 932 |
| **MW [kDa]** | 106.6 |
| **Score MS Amanda 2.0: MS Amanda 2.0** | 6258.8 |
| **Score Sequest HT: Sequest HT** | 84.78 |
| **Peptides (by Search Engine): MS Amanda 2.0** | 38 |
| **Peptides (by Search Engine): Sequest HT** | 19 |
| **Found in Sample:** | High |
